# SPACE-GM: geometric deep learning of disease-associated microenvironments from multiplex spatial protein profiles

**DOI:** 10.1101/2022.05.12.491707

**Authors:** Zhenqin Wu, Alexandro E. Trevino, Eric Wu, Kyle Swanson, Honesty J. Kim, H. Blaize D’Angio, Ryan Preska, Gregory W. Charville, Piero D. Dalerba, Ann Marie Egloff, Ravindra Uppaluri, Umamaheswar Duvvuri, Aaron T. Mayer, James Zou

**Affiliations:** Enable Medicine, Menlo Park, CA 94025, USA; Department of Chemistry, Stanford University, Stanford, CA 94305, USA; Department of Electrical Engineering, Stanford University, Stanford, CA 94305, USA; Department of Computer Science, Stanford University, Stanford, CA 94305, USA; Department of Pathology, Stanford University, Stanford, CA 94305, USA; Department of Pathology and Cell Biology, Columbia University, New York, NY 10027, USA; Department of Surgery, Brigham and Women’s Hospital, Boston, MA 02115, USA; Department of Medical Oncology, Dana-Farber Cancer Institute, Boston, MA 02115, USA; Department of Otolaryngology, University of Pittsburgh, Pittsburgh, PA 15213, USA; Department of Biomedical Data Science, Stanford University, Stanford, CA 94305, USA

## Abstract

Multiplexed immunofluorescence imaging enables high-dimensional molecular profiling at subcellular resolution. However, learning disease-relevant cellular environments from these rich imaging data is an open challenge. We developed SPAtial CEllular Graphical Modeling (SPACE-GM), a geometric deep learning framework that flexibly models tumor microenvironments (TMEs) as cellular graphs. We applied SPACE-GM to 658 head-and-neck and colorectal human cancer samples assayed with 40-plex immunofluorescence imaging to identify spatial motifs associated with cancer recurrence and patient survival after immunotherapy. SPACE-GM is substantially more accurate in predicting patient outcomes than previous approaches for modeling spatial data using neighborhood cell-type compositions. Computational interpretation of the disease-relevant microenvironments identified by SPACE-GM generates insights into the effect of spatial dispersion of tumor cells and granulocytes on patient prognosis.

## Introduction

Tumor microenvironments (TMEs) are complex niches characterized by cellular, molecular, and genetic heterogeneity. Current research and clinical practice have begun to reflect this complexity, with studies atlasing diseased cells in an unbiased fashion^1,2^ and novel therapies increasingly targeted to non-cancer cells, including immune and stromal compartments^3^. Just as the functions of healthy tissues depend on the spatial organization of cells, tumor pathology may depend on the spatial organization of the TME^4^.

In situ molecular profiling techniques, including spatial transcriptomic^5–7^ and proteomic^8–10^ techniques, increasingly enable high-dimensional, high-resolution characterization of TMEs and other tissues. Co-detection by indexing^8^ (CODEX) is an in situ molecular profiling technique based on iterative hybridization and fluorescence imaging of DNA-barcoded antibodies, enabling multiplexed quantification of 40 or more antigens from histological specimens at subcellular resolution.

While these new spatial technologies capture rich cellular and neighborhood information, analysis of the spatial data presents new challenges. In particular, how to identify biologically meaningful microenvironments from the rich spatial data and how to characterize the disease relevance of microenvironments are important open questions.

Previous works typically assign cells to cellular neighborhoods according to the cell-type compositions of their immediate neighbors^8,11–13^. However, these approaches may miss local spatial relationships between cells. Moreover, since the neighborhoods are generated in a purely unsupervised fashion, they have limited insight on which microenvironments are disease-relevant. We hypothesize that local spatial arrangements of cells beyond composition could encode rich disease-relevant information.

Here, we present SPAtial CEllular Graphical Modeling (SPACE-GM), a geometric deep learning framework that employs a graph neural network (GNN) to flexibly model cellular niche structures, or microenvironments, as subgraphs. Each node of the subgraph corresponds to a cell represented by its multiplexed protein levels and the edges capture neighbor relations. We apply SPACE-GM to three clinically annotated CODEX datasets and show that it identifies disease-relevant microenvironments that accurately predict patient-level phenotypes. We show that SPACE-GM generalizes across studies and disease contexts. Moreover, by analyzing the network embeddings, we derive specific insights on how local structural dispersion explains patient prognosis and treatment response.

There has been increased interest in applying graph-based deep learning methods to spatial cellular structures in recent literature^14–16^. Graph neural networks^17,18^ (GNNs), a class of deep learning methods designed for graph structures, have been applied to a variety of analysis tasks, including cell type prediction^19^, representation learning^20^, cellular communication modeling^21^ and tissue structure detection^22^. As most of these methods are designed for cellular property modeling, there still exists a gap between cellular-level graph analysis and patient-level phenotypes. SPACE-GM bridges this gap by training models using microenvironments as inputs to predict patient phenotypes. Interpretation of SPACE-GM sheds light on how cellular spatial arrangements impact disease and treatment outcomes.

## Results

### Geometric deep learning models cellular microenvironments

To model cellular communities, we first develop a pipeline to segment and classify individual cells from CODEX data (Methods). We then infer the 2D spatial structure of cells by constructing a Delauney triangulation and Voronoi diagram of cell centroid coordinates (Fig. 1A, middle panel). Lastly, we transform the data into a graph, defining cells as nodes and Delauney neighbors as edges (Fig. 1A, right panel).

**Figure 1.**
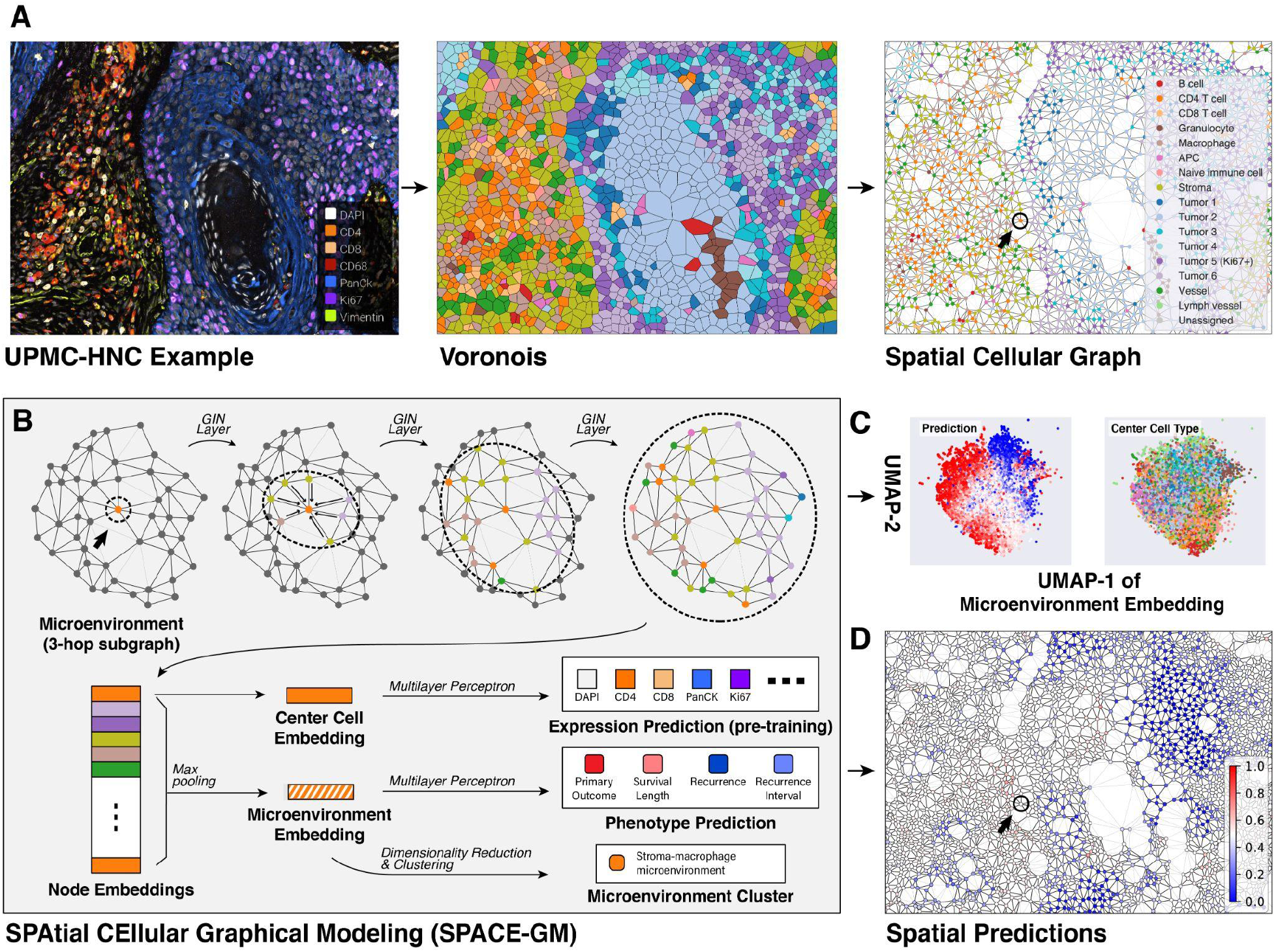
Geometric deep learning on multiplexed immunofluorescence imaging. **(A)** Pre-processing of CODEX data: we first transform the multiplexed fluorescence imaging data into Voronoi diagrams. Then we construct spatial cellular graphs from the Voronoi polygons, where each node is one cell and edges indicate adjacent cells. **(B)** Model structure of SPACE-GM: on the query node (marked by arrow and circle), a 3-layer GIN is applied to read the structure of its 3-hop neighborhood, which we call a microenvironment. Embeddings from the GIN are then used to predict cellular and phenotypic properties. **(C)** Dimensionality reduction (UMAP) of microenvironment embeddings from SPACE-GM. The left panel is colored by model prediction on the microenvironment, the right panel is colored by the center cell type. **(D)** Predictions on microenvironments are aggregated over the whole CODEX sample.

With the graphical representation as input, we propose SPAtial CEllular Graphical Modeling (SPACE-GM), a geometric deep learning tool that reads spatial cellular community structures in TMEs. SPACE-GM employs a graph isomorphism network (GIN)^23^ as the backbone and multiple multilayer perceptrons (MLP) as prediction heads (Methods).

SPACE-GM treats cellular graphs of TMEs as collections of local subgraphs: center nodes (cells) and their n-hop spatial neighbors (i.e., nodes within a graph distance of n edges from the center node). Empirically, we find that 3-hop neighborhoods–corresponding to 40 cells on average–is a suitable choice where the size of microenvironments and model performance is balanced (Supplementary Notes and Supplementary Fig. 4). We hereafter refer to these 3-hop local subgraphs as “microenvironments” (Fig. 1B). Correspondingly, a 3-layer GIN is employed in SPACE-GM, which constructs embeddings for center cells and microenvironments based on node features (e.g., one-hot encoded cell types) and edges (graph connectivity). Embeddings can then be passed through the prediction heads to generate estimates for cellular properties (e.g., center cell expression profile) and patient-level phenotypic properties (e.g., survival outcome of the patient).

In practice, we first pre-train SPACE-GM on cellular property prediction tasks, then finetune the backbone for patient phenotype predictions, both with microenvironment inputs (Fig. 1B, bottom row). In the text below, “SPACE-GM” and “SPACE-GM no-pretraining” represent models trained with or without the pre-training stage. During inference, we collect predictions from all individual microenvironments from test samples and perform mean-aggregation to derive patient-level predictions (Methods and Supplementary Notes).

### Applying SPACE-GM to head and neck cancer and colorectal cancer

To demonstrate the ability of SPACE-GM to model biologically and clinically relevant signals, we generated three 40-plex CODEX datasets from primary human cancer biopsies. Tissues were collected at Stanford University, University of Pittsburgh Medical Center (UPMC) and Dana Farber Cancer Institute (DFCI). In total, 658 samples were imaged, representing 139 head and neck cancer^24^ (HNC) patients and 110 colorectal cancer^25^ (CRC) patients (Fig. 2A). We refer to these datasets as UPMC-HNC, Stanford-CRC, and DFCI-HNC. Samples were annotated with clinical data, including patient survival, disease recurrence, and response to therapy (Supplementary Table 1).

**Figure 2.**
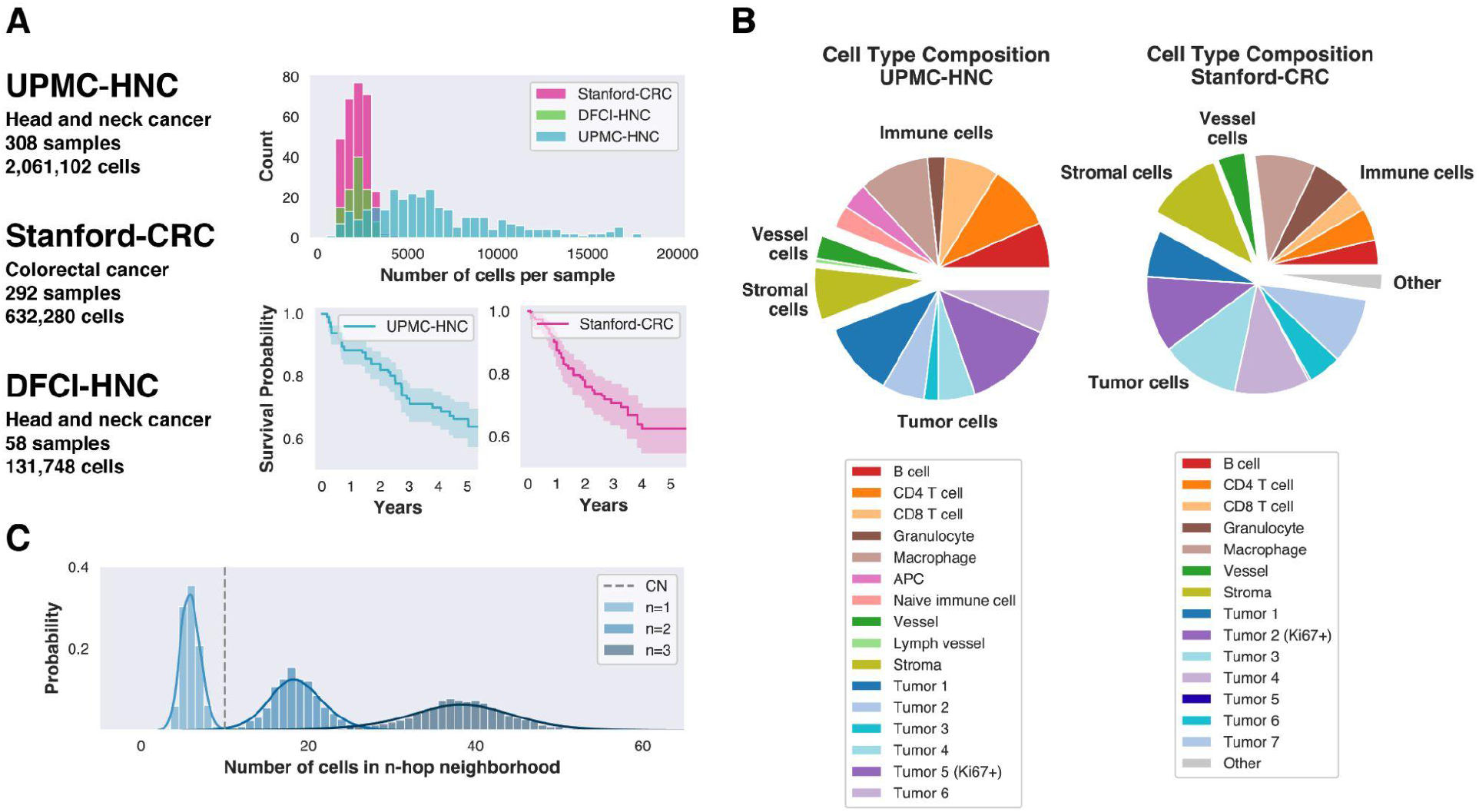
CODEX sample and cell characteristics. **(A)** Distributions of the number of cells per sample and overall survival curves. **(B)** Cell type compositions of UPMC-HNC and Stanford-CRC datasets. See Supplementary Fig. 1 for composition of DFCI-HNC dataset. **(C)** Size of n-hop neighborhoods (n=1, 2, 3) in UPMC-HNC. SPACE-GM utilizes a 3-layer GIN to read 3-hop neighborhoods (microenvironment). Dashed line indicates the neighborhood size (10 cells) commonly used in previous works^12^. SPACE-GM captures substantially larger neighborhoods.

CODEX samples were transformed into graphical representations following the pipeline described above. We extracted microenvironments by enumerating subgraphs of 3-hop neighbors within a distance threshold of 75 μm around each cell. The median microenvironment contains 38 cells, larger than previously proposed cellular neighborhoods^8,12^ (Fig. 2C). SPACE-GM is trained following the pre-training and/or finetuning approach, in which protein expression of the center cell is used as the pre-training task, and clinical annotations are used as phenotype prediction labels.

### SPACE-GM predicts patient phenotypes from cell microenvironments

We apply SPACE-GM to predict survival and recurrence outcomes for UPMC-HNC (Table 1) and Stanford-CRC (Table 2) patients. For each prediction task, SPACE-GM is trained with around 70% of the samples and tested on the remaining unseen samples from different coverslips (Methods). SPACE-GM achieves good performance on both datasets, with area under the curve (ROC-AUC) above 0.85 on all binary classification tasks and concordance index (C-index) around 0.8 on the survival analysis task in UPMC-HNC. Stanford-CRC shows slightly worse performance, likely due to having fewer samples and cells.

**Table 1.** Prediction performance on UPMC-HNC and DFCI-HNC tasks. Binary classification and hazards columns report average performances on the validation folds of UPMC-HNC. The generalization column reports the cross-study (UPMC-HNC to DFCI-HNC) prediction performance.

| Model | Binary Classification (ROC-AUC) |  |  | Hazards Model (C-index) | Generalization (ROC-AUC) |
| --- | --- | --- | --- | --- | --- |
|  | Primary Outcome | Recurrence | HPV Infection | Survival Length | Primary Outcome<sup>1</sup> |
| Linear on composition (sample) | 0.783 | 0.852 | 0.870 | 0.696 | 0.731 |
| MLP on composition (sample) | 0.771 | 0.869 | 0.879 | 0.721 | 0.754 |
| Linear on composition (microenvironment) | 0.774 | 0.823 | 0.864 | 0.700 | 0.799 |
| MLP on composition (microenvironment) | 0.814 | 0.832 | 0.891 | 0.751 | 0.806 |
| SPACE-GM no-pretraining | 0.854 | 0.882 | 0.918 | 0.778 | 0.853 |
| SPACE-GM | **0.867** | **0.883** | **0.926** | **0.799** | **0.873** |

<sup>1</sup>Models trained with UPMC-HNC primary outcome tasks are applied to DFCI-HNC samples, predictions are evaluated for the binary primary tumor response task.

**Table 2.** Prediction performances on Stanford-CRC tasks. All columns report average performances on the validation folds of Stanford-CRC.

| Model | Binary Classification (ROC-AUC) |  | Hazards Model (C-index) |  |
| --- | --- | --- | --- | --- |
|  | Primary Outcome | Recurrence | Survival Length | Recurrence Interval |
| Linear on composition (sample) | 0.563 | 0.592 | 0.524 | 0.537 |
| MLP on composition (sample) | 0.551 | 0.542 | 0.577 | 0.570 |
| Linear on composition (microenvironment) | 0.576 | 0.599 | 0.562 | 0.569 |
| MLP on composition (microenvironment) | 0.547 | 0.491 | 0.525 | 0.519 |
| SPACE-GM no-pretraining | 0.684 | 0.675 | 0.642 | 0.669 |
| SPACE-GM | **0.739** | **0.696** | **0.655** | **0.713** |

For context, we compare SPACE-GM’s clinical prediction performances against alternative composition-based methods, applied on either whole samples or subgraphs. As input to these baseline models, we use whole graphs or the same subgraphs of 3-hop neighborhoods (microenvironment), but featurized as cell-type composition vectors (Methods). Compared with graph representations, composition vectors collapse spatial structure, losing information about the relative spatial arrangement of cells within a subgraph. Both a linear model (logistic regression or proportional hazards regression) and a multilayer perceptron are trained and evaluated following the same pipeline (Methods).

SPACE-GM consistently outperforms baseline methods on both classification and hazards modeling (time-to-event) tasks (Table 1 and Supplementary Table 2). Uncertainty of the performance metrics is calculated by bootstrapping (Supplementary Table 3 and Supplementary Notes), which demonstrates consistent advantages from SPACE-GM. Model pre-training confers an additional advantage on all prediction tasks. On the most challenging dataset Stanford-CRC, where composition-based methods generate nearly random predictions, SPACE-GM demonstrates a robust test set performance (Table 2).

On a representative task - primary outcome of the UPMC-HNC dataset - we plot receiver operating characteristic (ROC) curves for one of the test folds and observe a substantial advantage for SPACE-GM predictions over baseline methods (Fig. 3A). It is also worth noting that MLP based on microenvironment compositions outperforms the same model using whole graph compositions, which is also reflected in most of the other prediction tasks.

**Figure 3.**
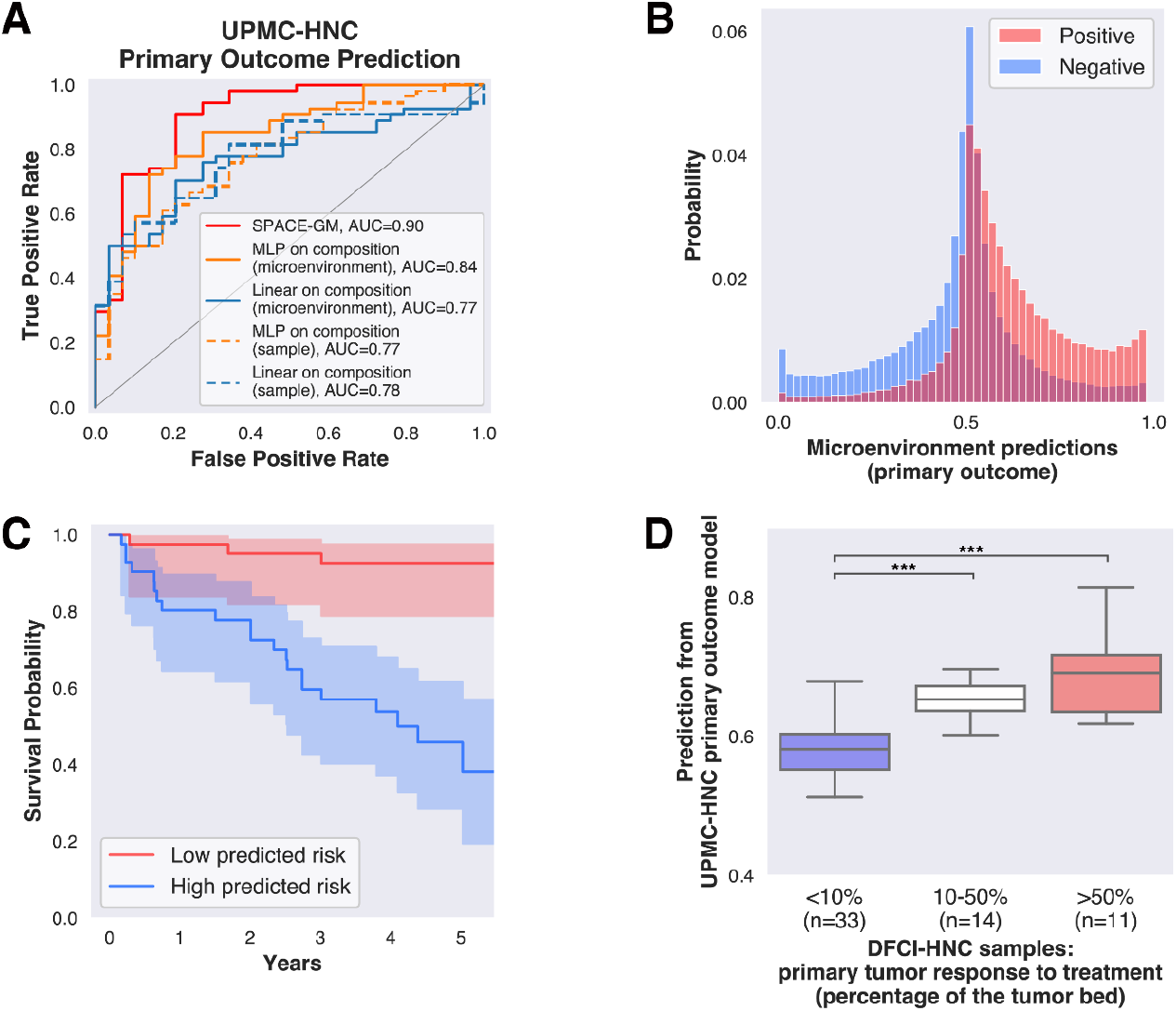
Model predictions on primary outcome and survival length in HNC. **(A)** Receiver operating characteristic (ROC) curves of different methods on a representative task: primary outcome prediction in the UPMC-HNC dataset. SPACE-GM outperforms all other methods by a comfortable margin. One of the two validation folds is shown here, results are representative of the other fold. **(B)** Distribution of microenvironment prediction values on the test set from SPACE-GM. The majority of the predictions are neutral, with a small portion of positive/negative predictions. **(C)** Survival curves of patients in the test set grouped by SPACE-GM predicted risk. Two groups are separated by the median predicted risk. Survival probability of the low-risk group is significantly higher than the high-risk group (*P* < .001, log-rank test). **(D)** Predictions on the DFCI-HNC dataset from the primary outcome model trained with UPMC-HNC samples. SPACE-GM predicts significantly higher scores (*P* < .001, two-sample t-test) for the high response group (red box and white box) than for the low response group (blue box).

The observation above indicates that the superiority of geometric deep learning stems in part from our microenvironment-aggregation strategy. Fig. 3B plots the histogram of individual microenvironment predictions before aggregation. In addition to the difference in distributions of prediction values between positive and negative samples, we also notice that the majority of predictions, regardless of label, are neutral. This suggests that most spatial cellular structures are shared among different patients, while only a small fraction of motifs are highly indicative of patient outcomes. Identifying and characterizing these disease-relevant microenvironments will be highly informative for better diagnostics and therapeutics.

Fig. 3C plots the survival curves of two patient cohorts in the test data, separated by median predicted risk from SPACE-GM (trained under the survival length task). We observe a significant difference (*P* < .001, log-rank test) in survival probability between the low-risk group and the high-risk group. The survival curves stratitifed by SPACE-GM risk scores show greater separation than curves stratified using cell-type composition risk scores (Supplementary Fig. 3), suggesting that SPACE-GM captures more granular spatial motifs predictive of patient mortality.

### Cross-study generalizability of SPACE-GM

We next seek to evaluate the generalizability of SPACE-GM across datasets. The DFCI-HNC contains CODEX samples collected both before and after neoadjuvant therapy in 29 patients. Though patient survival outcomes in this cohort are not available, samples are annotated based on the degree of pathologic tumor response (PTR) in surgically resected tumors after neoadjuvant anti-PD1 therapy^26^, which serves as a proxy for primary outcome in this experiment.

In the experiment, we train models on primary outcome labels from the UPMC-HNC dataset with integrated cell types (Supplementary Notes) and directly apply them to pre-therapy DFCI-HNC samples to predict therapeutic response. Despite potential batch effects resulting from tissue handling, biopsy size (Fig. 2A), and independently-generated cell labels (Methods), we find that SPACE-GM generates robust estimations, with accuracy surpassing all baseline composition-based models (Table 1).

To examine this result in greater detail, we group SPACE-GM predictions by pathologist-annotated PTR categories (Fig. 3D) and find that, strikingly, model predictions align well with fine-grained PTR categories. A two-sample t-test suggests that there is significant differences in predictions between <10% and >10% responder samples (*P* < .001, two-sample t-test). These results demonstrate the robustness and generalizability of SPACE-GM.

### Defining disease-relevant cell microenvironments with SPACE-GM

Motivated by the superior performance of SPACE-GM over composition-based baselines methods, we further investigate how characteristics of cellular community structures beyond composition are predictive of clinical outcomes. The prediction accuracy of SPACE-GM suggests that its embedding space, which learns to present each microenvironment by a numerical vector, is informative of the phenotypes of interest. Visualization with UMAP shows that patient phenotypes are well-separated in the embedding space (Fig. 1C and Supplementary Fig. 6).

We use the primary outcome task from UPMC-HNC as an example to demonstrate how to interpret SPACE-GM embeddings to generate biological hypotheses related to patient outcomes. We first cluster all the microenvironments based on SPACE-GM embeddings (Methods). Clusters show distinct cell type enrichment patterns and prediction values (Fig. 4A).

**Figure 4.**
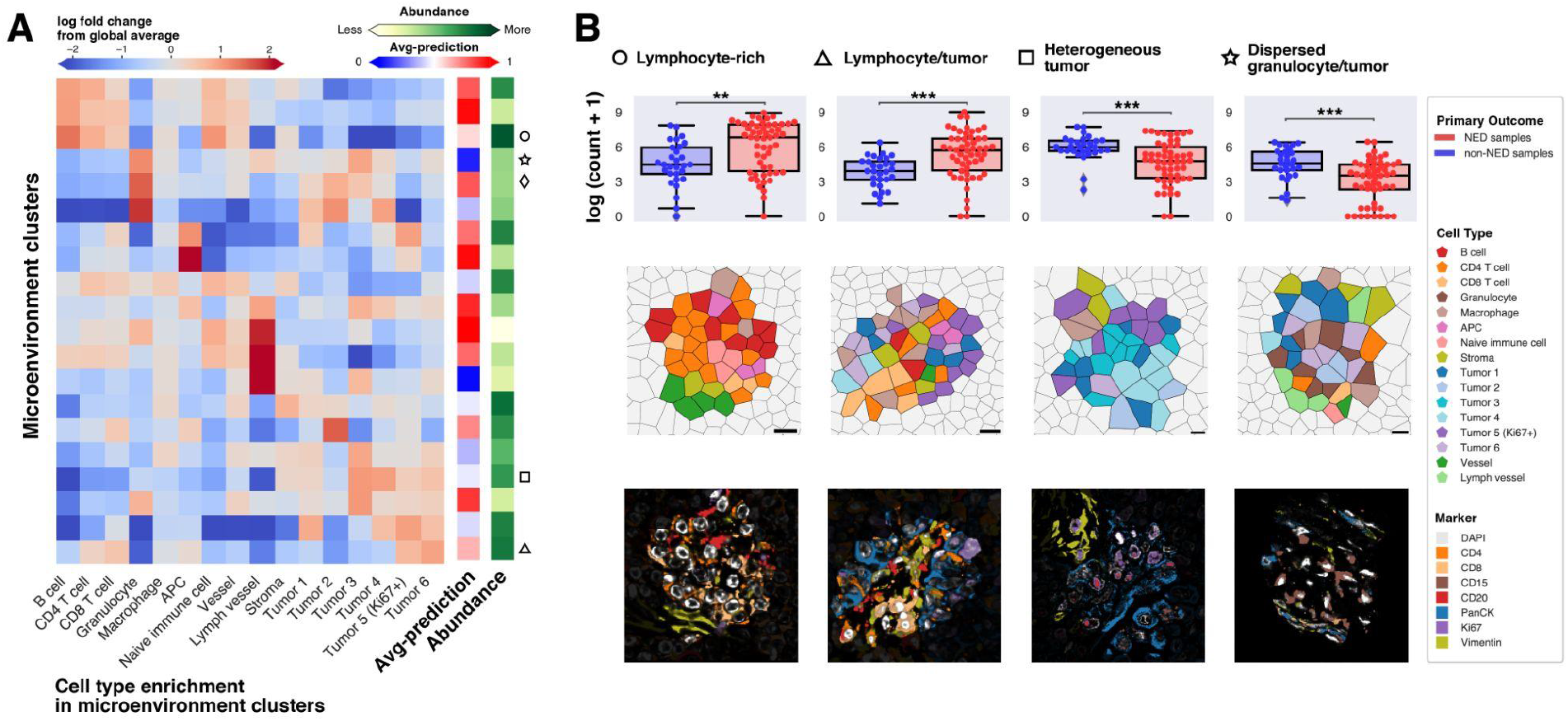
Clustering of SPACE-GM embeddings identifies disease-relevant microenvironments. **(A)** 20 clusters are identified by fitting a K-means clustering (K=20) on microenvironment embeddings. Clusters have different enrichment for cell types and SPACE-GM predictions. **(B)** (top row) On four representative clusters, we count the appearance of corresponding microenvironments in test samples, which show significant differences between NED and non-NED samples. (middle row) Voronoi diagrams and (bottom row) raw CODEX images of sample microenvironments from these clusters are illustrated.

Four clusters of interest are displayed in Fig. 4B, all of which show significant differences in frequency between positive (no evidence of disease, NED) and negative (non-NED) patient samples, indicating that they possess informative cellular motifs (Fig. 4B, *P* = .002 for lymphocyte-rich microenvironments, *P* < .001 for the rest, two-sample t-test).

Voronoi diagrams and raw multiplexed fluorescence images are visualized for example microenvironments in each cluster (Fig. 4B middle and bottom row). We name the clusters based on their cell type compositions and spatial cellular structure patterns: a lymphocyte-rich cluster, indicating high immune activity (circle symbol); a mixed lymphocyte/tumor cell cluster (triangle); a heterogeneous tumor cluster characterized by various subtypes of tumor cells (square); and a tumor cell and myeloid cell (granulocytes and macrophages) cluster characterized by dispersed spatial distributions (star). The circle and triangle clusters appear more in positive outcome samples and are thus associated with better prognosis. In contrast, the square and star clusters are more enriched in negative outcome patients.

### In silico permutations suggest disease-relevant spatial motifs in microenvironments

Clusters of disease-relevant microenvironments indicated correlations between microenvironment structures and clinical phenotypes, in which we are especially interested in structural characteristics beyond composition. For example, the formation of distinctly-shaped boundaries between immune and tumor cells could indicate distinct pathways and programs related to tumor progress or immune control.

To directly test how structural arrangements impact microenvironment function, we computationally permute nodes in microenvironments and measure SPACE-GM outcome predictions on the permuted graphs. Our permutations are designed to vary the amount of dispersion allowed in the final graph. Thus, a “dispersed permutation” in our scheme rearranges all cells amongst each other, resulting in highly mixed graphs. A “coherent permutation” extracts target cell types, and places them into sectors around the center cell, resulting in cells of the same type appearing together (Methods). In this section, we pick two representative clusters discovered above and evaluate how permutations towards the two extremes of dispersion levels affect model predictions.

In the heterogeneous tumor cluster, we observe a group of microenvironments enriched in different tumor subtypes being highly indicative of non-NED outcomes. We compare its appearance in all samples against each of the individual tumor subtypes and stromal cells (Fig. 5A). Interestingly, no individual tumor subtype is differentially enriched between positive and negative samples.

**Fig. 5.**
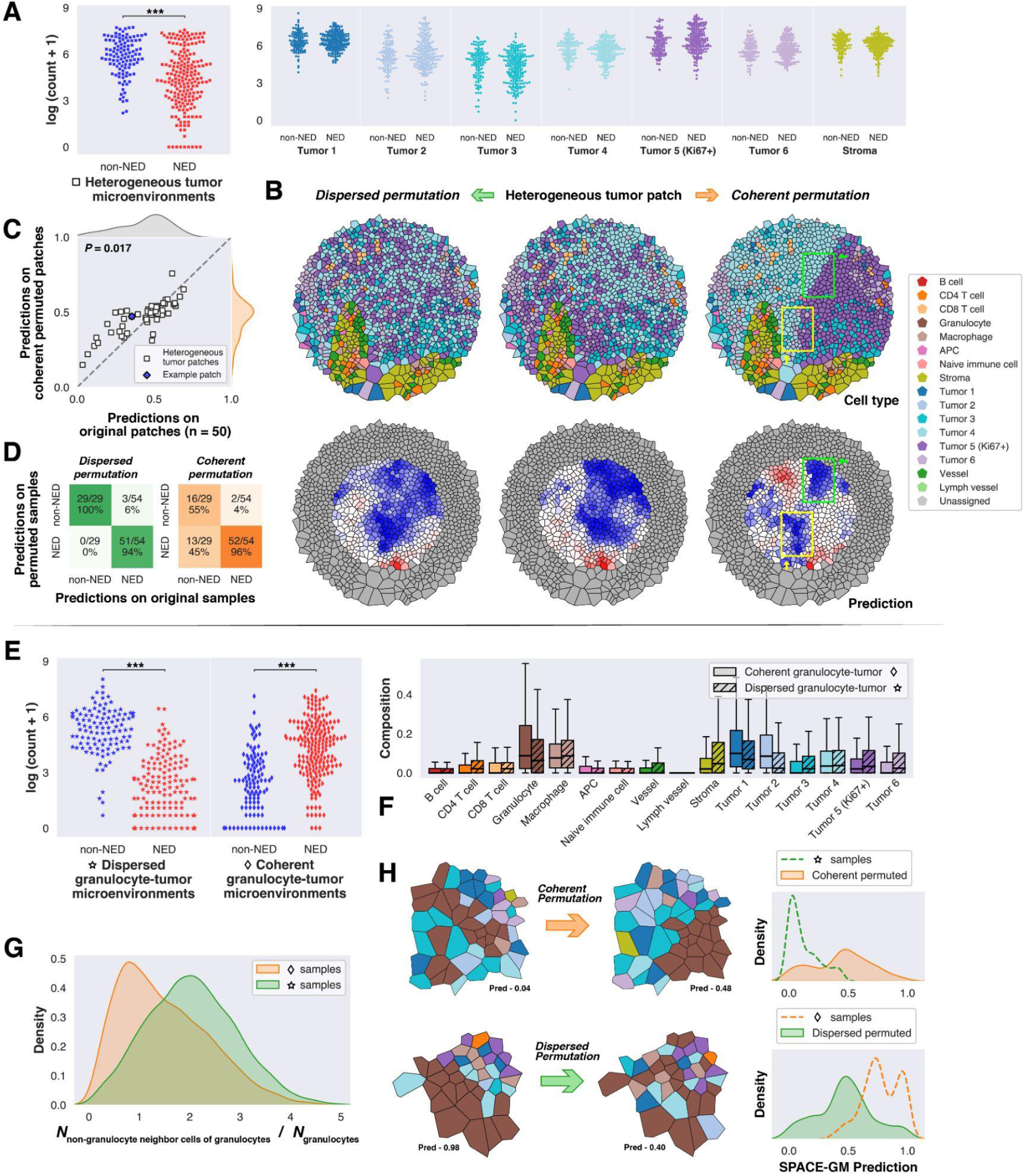
Permutation of nodes in microenvironments helps identify cell-cell interaction affecting predictions. **(A)** Swarm plots show counts per sample of heterogeneous tumor microenvironments and each individual tumor subtype. Note that even though no specific tumor subtype shows a strong correlation with phenotype, heterogeneous tumor microenvironments display a significant difference between NED and non-NED samples (*P* < .001, two-sample t-test). **(B)** Example heterogeneous tumor patch (middle column) and its two permuted versions (left and right column) are displayed as Voronoi diagrams. Note that in the coherent permuted patch, areas of negative prediction values (blue) overlap with tumor subtype boundaries. **(C)** Scatter plot of predictions on 50 heterogeneous tumor patches. Coherent permuted patches have significantly higher predictions than their original patches (*P* = .02, Wilcoxon signed-rank test). **(D)** Confusion matrices of predicted classes between original samples and permuted samples. 45% of the samples originally classified as non-NED are predicted as NED after coherent permutation. **(E)** Swarm plots of two granulocyte and tumor rich microenvironments: distributions in NED and non-NED samples show opposite trends. **(F)** Cell type compositions of the two granulocyte-tumor microenvironments are similar. **(G)** A heuristic measure: count of non-granulocyte neighbors of granulocytes divided by count of granulocytes, is calculated for the two microenvironment groups, distributions show significant differences (*P* < .001, two-sample t-test). **(H)** Example microenvironments from the two groups undergoing permutation are displayed as Voronoi diagrams. Distributions (Kernel Density Estimate) of SPACE-GM predictions on selected microenvironments and their permuted versions are plotted in the right column, note the shift of distributions due to permutations.

To test whether the arrangement of tumor subtypes confers the prediction bias on these samples, we select regional patches (Methods) and perform the permutations described above. Fig. 5B illustrates an example patch (middle column) and the two permuted versions (left and right columns) that mix/separate *Tumor 4* and *Tumor 5 (Ki67+)*. No major change is observed after dispersed permutation as the original patch is highly mixed. In contrast, when subject to coherent permutation, mean-aggregated prediction increases. Moreover, we observe that the boundaries of different tumor subtypes (yellow and green boxes) overlap with the negative prediction regions, suggesting that the mixing of these tumor subtypes explains the negative prediction in this example.

We support this result on a diverse set of 50 heterogeneous tumor regional patches (Fig. 5C and Methods): SPACE-GM predictions on coherent permuted patches are significantly higher (*P* = .02, Wilcoxon signed-rank test). This trend could be even extended to the whole-sample level. On all of the 83 samples from the test set, we run coherent permutations and evaluate changes in predictions (Fig. 5D). Among the 29 samples that are predicted to have negative outcomes, 45% of them have predictions altered after coherent permutation, compared with 0% after dispersed permutation. These results suggest that the spatial mixing of tumor subtypes is a negative predictor of patient outcomes captured by SPACE-GM.

We next evaluate the spatial organization of two SPACE-GM embedding clusters, termed dispersed and coherent granulocyte-tumor microenvironments, with similar cell-type compositions (Fig. 5F) but opposite outcome enrichment (Fig. 5E). Visualization of examples from the two clusters (Fig. 5H, left column) suggests that the difference could potentially be attributed to the dispersion of granulocytes, and we verified this by calculating the number of non-granulocyte neighbors for 10,000 granulocytes from these two clusters (Fig. 5G). The distribution shows a significant right shift (*P* < .001, two-sample t-test) for dispersed granulocyte/tumor cluster samples, indicating that granulocytes are indeed more dispersed in this cluster.

To further characterize the relations between the dispersion of granulocytes and predictions, we perform the two types of permutations (Methods) on corresponding microenvironments to reverse the spatial organization patterns of granulocytes (Fig. 5H, middle column and Supplementary Fig. 6). Strikingly, coherent permutation of granulocytes in dispersed clusters results in improved predicted prognoses. Conversely, dispersed permutation of coherent granulocyte clusters results in microenvironments with poorer prognoses (Fig. 5H, right column).

## Discussion

In this work, we present SPACE-GM, a graph neural network model that predicts clinical properties of patient tumor samples with multiplexed immunofluorescence inputs. We evaluate prediction performances on three independently collected patient cohorts and demonstrate the predictive superiority of SPACE-GM over composition-based methods, suggesting that spatial cellular arrangements beyond composition could encode phenotype-related information. We further conduct dimensionality reduction and clustering on the microenvironment embeddings from SPACE-GM to identify discrete groups of disease-relevant microenvironments. Permutation on two example clusters unveils the relations between spatial dispersion of certain cell types and patient outcomes.

More specifically, SPACE-GM learns that the dispersion of molecularly distinct tumor subtypes could have a negative impact on patient survival outcomes. Of note, similar correlations between cell type mixing (or tumor subtype mixing) and poorer outcomes have been observed in recent literature^27,28^. In another experiment, we verified that the dispersion level of granulocytes within the TME is relevant to SPACE-GM predictions of primary outcomes. Interestingly, similar trends have been reported in both head and neck cancer^29,30^ and other solid cancers^31^, in which multiple molecular pathways explaining how neutrophils promote tumor growth^32,33^ and metastasis^34,35^ are presented.

Results discussed in this work demonstrate the value of modeling patient phenotypes with graph-based microenvironment inputs, from which connections between cellular community structures and patient-level phenotypes can be established and validated. However, we acknowledge that certain caveats still exist in our framework: Inference and analysis of unseen data are still complicated due to the limitations of cell segmentation and classification approaches. The lack of a unified image-cell type dictionary hinders the generalization of trained models on new datasets or unseen cell types. Comprehensive results from human cell consortia efforts^36,37^ as well as computational methods that accommodate unseen cell types^19^ could potentially be incorporated to overcome these limitations.

Additionally, dense per-cell predictions by SPACE-GM are mean-aggregated to maximize prediction performance in this work, based on the benchmarking results of basic aggregation methods. As illustrated in Fig. 1D and Fig. 3B, highly disease-relevant microenvironments are usually less prevalent, the spatial distribution of which in a sample could potentially contain information about tissue architecture at different spatial scales. We foresee that more sophisticated techniques (e.g., hierarchical aggregation on different spatial scales) could be employed at this step, revealing insights into the interplay between tissue-level architectures and patient-level phenotypic properties.

SPACE-GM is a versatile framework to capture disease-relevant motifs from microenvironments. We applied SPACE-GM to analyze CODEX data in this work. The framework of using local graphs to characterize disease-relevant spatial motifs can be extended to other measurement modalities such as spatial transcriptomics, and this is an interesting direction of future work. Disease-relevant microenvironment embeddings and dense predictions of target phenotypes could be further coupled with downstream analysis (e.g., permutation) to reveal relationships between cellular community structure and patient-level phenotypes.

## Methods

### CODEX Data Collection

All samples are prepared, stained, and acquired following CODEX User Manual Rev C (https://www.akoyabio.com).

#### Coverslip preparation

Coverslips are coated with 0.1% poly-L-lysine solution to enhance adherence of tissue sections prior to mounting. The prepared coverslips are washed and stored according to the guidelines in the CODEX User Manual.

#### Tissue sectioning

formaldehyde-fixed paraffin-embedded (FFPE) samples are sectioned at a thickness of 3-5 μm on the poly-L-lysine coated glass coverslips.

#### Antibody conjugation

Custom conjugated antibodies are prepared using the CODEX Conjugation Kit, which include the following steps: (1) the antibody is partially reduced to expose thiol ends of the antibody heavy chains; (2) the reduced antibody is conjugated with a CODEX barcode; (3) the conjugated antibody is purified; (4) Antibody Storage Solution is added for antibody stabilization for long term storage. Post-conjugated antibodies are validated by SDS-polyacrylamide gel electrophoresis (SDS-PAGE) and quality control (QC) tissue testing, where immunofluorescence images are stained and acquired following standard CODEX protocols, then evaluated by immunologists.

#### Staining

CODEX multiplexed immunofluorescence imaging was performed on FFPE patient biopsies using the Akoya Biosciences PhenoCycler platform (also known as CODEX). 5 μm thick sections were mounted onto poly-L-lysine-treated glass coverslips as tumor microarrays. Samples were pre-treated by heating on a 55 °C hot plate for 25 minutes and cooled for 5 minutes. Each coverslip was hydrated using an ethanol series: two washes in HistoChoice Clearing Agent, two in 100% ethanol, one wash each in 90%, 70%, 50%, and 30% ethanol solutions, and two washes in deionized water (ddH2O). Next, antigen retrieval was performed by immersing coverslips in Tris-EDTA pH 9.0 and incubating in a pressure cooker for 20 minutes on the High setting, followed by 7 minutes to cool. Coverslips were washed twice for two minutes each in ddH2O, then washed in Hydration Buffer (Akoya Biosciences) twice for two minutes each. Next, coverslips were equilibrated in Staining Buffer (Akoya Biosciences) for 30 minutes. The conjugated antibody cocktail solution in Staining Buffer was added to coverslips in a humidity chamber and incubated for 3 hours at room temperature or 16 hours at 4 °C. After incubation, the sample coverslips are washed and fixed following the CODEX User Manual.

#### Data acquisition

Sample coverslips are mounted on a microscope stage. Images are acquired using a Keyence microscope that is configured to the PhenoCycler Instrument at a 20X objective. All of the sample collections were approved by institutional review boards.

### Cell segmentation and classification

After image preprocessing, we applied a neural network-based cell segmentation tool DeepCell^38^ on DAPI image channels to identify nuclei, and these nuclear masks were dilated to obtain whole-cell segmented cells. Nuclear segmentation masks were stochastically dilated by flipping pixels with a probability equal to the fraction of positive neighboring pixels. This dilation was repeated for 9 cycles for all CODEX data.

On each CODEX sample, given the segmentation of individual cells, single cell expression was computed for biomarker *j* with the following steps^39^:

- Compute the mean expression value across pixels within the cell segmentation mask. Denote the mean expression value of cell *i* as 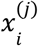, and denote the array of all expression values 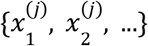 as *X*^(*j*)^.
- Normalize the expression value using quantile normalization and arcsinh transformation:

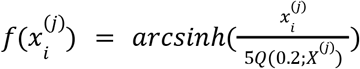

in which *Q*(0. 2; *X*^(*j*)^) represents the 20-th quantile of *X*^(*j*)^ and *arcsinh* is the inverse hyperbolic sine function. Denote the array of all normalized expression 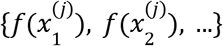 as *f*(*X*^(*j*)^).
- Calculate z-score of normalized expression value:

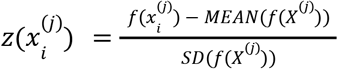

To classify cells, we first obtained a cell-by-marker expression matrix filled with preprocessed expression values 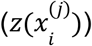, then a Principal Component Analysis (PCA) model was applied to extract the top 20 Principal Components (PCs). We constructed a k-nearest-neighbor graph (kNN, k = 30) on the top 20 PCs of the expression matrix, then performed Louvain graph clustering^40^ on the result. Clusters were manually annotated according to their cell biomarker expression patterns. This procedure was performed on a subset of 10,000 cells and subsequently used to train a KNN algorithm to predict cell type from the normalized expression vector. This algorithm was used to transfer labels to the entire dataset. The average expression of each cell type for all the three datasets used in this work is visualized in Supplementary Fig. 2.

### Construction of spatial cellular graphs and microenvironments

For each multiplexed fluorescence image, we identified individual cells by the segmentation and classification pipeline stated above. The set of cells will be represented by a set of discrete points located at cellular centroids. 2D coordinates of these cellular centroids were determined by the segmentation masks of the corresponding cell nuclei.

To capture the spatial neighborhood relations, we run a Delauney triangulation operation on all the cellular centroids. Corresponding Voronoi diagrams can be uniquely determined by connecting the centers of the circumcircles. We employed the function `voronoi_regions_from_coords` from package `geovoronoi`^41^ in this step.

The graphical representation of multiplexed fluorescence images can then be determined by defining cellular centroids as nodes and neighboring Voronoi polygons (or edges in the Delauney triangulation) as edges. We further defined two types of edges based on the distance between cellular centroids: edges shorter than 20 μm are treated as neighboring edges; edges longer than 20 μm are treated as distant edges. Edge types will be considered in the following neural network forward pass.

Microenvironment represents the local environment around a cell in the entire cellular community. Given the graphical representation derived above, we defined the microenvironment of a query cell as its n-hop neighborhood. Here the n-hop neighborhood includes all cells within a graph distance of n edges from the query cell. In this work, we applied n=3 in all datasets and added another constraint of physical distance: microenvironment of a query cell includes all cells that are in its 3-hop neighborhood and less than 75 μm away from the query cell.

### Data split and evaluation metrics

We evaluated SPACE-GM and other baseline methods on the UPMC-HNC and Stanford-CRC datasets. Samples were first split into training set and test set following the procedure below:

**UPMC-HNC** (Table 1, coverslip split)

- UPMC-HNC contains 308 samples collected from 7 batches/coverslips, class balance of clinical annotations are calculated for each coverslip.
- We proposed two validation folds:
  ○ Fold 1
    ■ 225 samples in the training set, 64% have positive primary outcomes;
    ■ 83 samples in the test set, 65% have positive primary outcomes;
  ○ Fold 2
    ■ 217 samples in the training set, 65% have positive primary outcomes;
    ■ 91 samples in the test set, 62% have positive primary outcomes;
  ○ In each fold, samples from 5 coverslips are used for training, samples from the remaining 2 coverslips are used for testing. Class balance of clinical annotations are kept similar between training and test sets.
  ○ Two validation folds have no overlapping test samples.
- Training / evaluation are run independently on the two validation folds, prediction performances for each task are averaged.

**UPMC-HNC** (Supplementary Table 2, patient cross validation)

- UPMC-HNC contains 308 samples collected from 81 patients.
- Patients are randomly split into four groups, samples are assigned to their corresponding patient groups.
- Training/evaluation following cross-validation scheme:
  ○ In each of the four independent runs, models are trained on samples from 3 patient groups and evaluated on samples from the remaining group.
  ○ Performances are averaged across the four runs.

**Stanford-CRC** (Table 2, coverslip split)

- Stanford-CRC contains 292 samples from 4 batches/coverslips, class balance of clinical annotations are calculated for each coverslip.
- We proposed two validation folds based on coverslip:
  ○ Fold 1
    ■ 229 samples in the training set, 75% have positive primary outcomes;
    ■ 63 samples in the test set, 71% have positive primary outcomes;
  ○ Fold 2
    ■ 220 samples in the training set, 78% have positive primary outcomes;
    ■ 72 samples in the test set, 64% have positive primary outcomes;
  ○ In each fold, samples from 3 coverslips are used for training, samples from the remaining coverslip are used for testing.
  ○ Two validation folds have no overlapping test samples or patients. Note that the two coverslips solely used for training are excluded from testing because of their different size/class balance.
- Training/evaluation are run independently on the two validation folds, and prediction performances for each task are averaged.

In this work, we use clinical annotations as prediction tasks. Annotations are categorized into two forms (Supplementary Table 1): binary classification (e.g., primary outcome) and hazards modeling (e.g., survival length). On binary classification tasks, we evaluate model performances by calculating area under the curve (AUC) of receiver operating characteristics (ROC); on hazards modeling tasks, we evaluate performances by calculating concordance index (C-index) between predicted hazards and observed events (recurrence or death).

### SPACE-GM and baseline methods

#### SPACE-GM

SPACE-GM consists of a Graph Isomorphism Network (GIN)^23^ backbone and multiple multilayer perceptron (MLP) prediction heads.

Inputs of SPACE-GM contain the local spatial graphical structures of microenvironments derived above, as well as identity and size of each cell in the microenvironments. More specifically, GIN has the following inputs:

- Node features
  ○ Cell type is formed as a one-hot vector of length *N_cell type_*, which is mapped to *N_cell type_* trainable embeddings of length 512 through a lookup table.
  ○ Other features: Cell size (in pixel) log-transformed and scaled to 0-1 range, a flag indicating if the cell is the center taking value 0 (if no) or 1 (if yes), are concatenated and transformed to a vector of length 512 through a trainable linear layer. Note that we experimented with explicitly adding normalized expression to node features, but the resulting models have more severe overfitting and lower test set performances. Expression is hence excluded from node features.
  ○ The two embeddings above are summed and used as the initial node embeddings for GIN. We denote the initial embedding of node *v* as 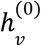.
- Edge features
  ○ Self-loop edges (connecting same nodes) are added to input graphs before forward pass.
  ○ Edges are divided into three classes: neighboring edge, distant edge, and self-loop edge. In each GIN layer, an edge is mapped to one of the three edge embeddings of length 512 through a lookup table. We denote the embedding of the edge between node *v* and node *u* in *k*-th layer as 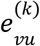, which is dependent on the edge type between *v* and *u*. Note that 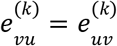.

SPACE-GM employed a 3-layer GIN, and in the *k*-th graph convolutional layer:

- Messages are calculated on each edge as:

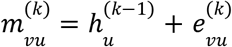 Note that edges in microenvironments are undirected, messages in both directions are calculated, though they will not necessarily be equal.
- Embedding of node is updated based on all incoming messages to *v*:

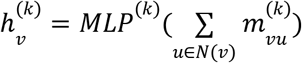

in which *N*(*v*) is the set of neighboring nodes of *v*, the self-loop edge guarantees *v* ∈ *N*(*v*), and *MLP*^(*k*)^ is the 2-layer MLP of the *k*-th layer.

Embeddings from the last graph convolutional layer are treated as node embeddings, in which the embedding of the center cell 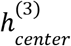 is used as input for expression prediction (pre-training of SPACE-GM). We aggregated node embeddings to generate the microenvironment embedding:

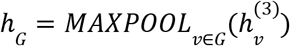

in which *G* represents the microenvironment, is a channel-wise maximum operation (`torch.nn.global_max_pool`). Microenvironment embeddings are used for sample phenotype predictions.

For expression and phenotype prediction tasks, we employed two separate 3-layer MLP with Leaky ReLU activation function, each with *N_task_* outputs, taking center cell embedding 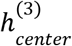 and microenvironment embedding *h_G_* as input respectively. For expression prediction tasks, we minimize squared L2 norm loss between predictions and labels (`torch.nn.MSELoss`). For binary classification tasks, we minimize binary cross entropy between sigmoid logit: outputs of the 3-layer MLP and class labels (`torch.nn.BCEWithLogitsLoss`). For hazards modeling tasks, we adapted the Cox partial likelihood for SGD introduced by Kvamme et al.^42^.

#### Baseline methods

Baseline methods in this work are constructed based on composition vector inputs. Composition vectors are calculated on whole-sample graphs or microenvironments. In both cases, we count the number of each cell type appearing in the graph/subgraph. The count vector of length *N_cell type_* is then normalized to the frequency vector, denoted as the composition vector.

We trained a linear model and a 3-layer MLP model on the composition vectors. Logistic regression and Cox regression are used for binary classification and hazards modeling tasks respectively as linear models. MLP used the same loss function as SPACE-GM introduced above.

#### Model training and inference

During training (of microenvironment-based methods), we randomly select microenvironments from all samples in the training set. Their labels came from center cells (expression prediction) or clinical annotations of the CODEX samples they belong to (phenotype prediction). Microenvironments are weighted based on their labels to balance loss of different classes. Adam optimizer^43^ is employed to minimize corresponding losses in different tasks.

SPACE-GM is first trained on the expression prediction task. After convergence, we retain the GIN backbone and connect it with the phenotype prediction head (initialized from scratch), and both modules are finetuned on the phenotype task. SPACE-GM no-pretraining has the same model structure as SPACE-GM, but all initialized from scratch. It is directly trained on the phenotype task until convergence; microenvironment-based MLP models follow the same training pipeline.

During inference, we run microenvironment-based models (SPACE-GM, MLP, etc.) on all microenvironments from the test set, generating dense per-node predictions for test samples. Predictions within the same CODEX sample are then mean-aggregated (microenvironment-aggregation, see Supplementary Notes and Supplementary Fig. 5 for discussion), the results of which are evaluated using corresponding metrics.

All models are implemented in Python `with scikit-learn`^44^, `pytorch`^45^, and `pytorch geometric`^46^. More details about the hyperparameters and implementations can be found in our github repository at https://gitlab.com/enable-medicine-public/space-gm.

### Clustering of microenvironment embeddings

To identify clusters of disease-relevant microenvironments, we applied dimensionality reduction and clustering on microenvironment embeddings. Clusters discussed in the main text are generated following the procedure below:

- 100,000 cells and their microenvironments from the training set are randomly sampled and extracted, denoted as the reference dataset;
- SPACE-GM (trained on the UPMC-HNC primary outcome task) is applied to all microenvironments in the reference dataset, their microenvironment embeddings (*h_G_*) and predictions are collected, denoted as reference embeddings and reference predictions;
- A Principal Component Analysis (PCA) model is initialized and fitted to the reference embeddings. We extracted the top 20 Principal Components (PCs), which captured >70% of total variance;
- A UMAP dimensionality reduction model^47^ is fitted on the top 20 PCs of reference embeddings, generating 2D visualizations of microenvironment space;
- A K-means clustering (K=20) is fitted on the top 20 PCs of reference embeddings to identify 20 clusters of microenvironments;

We then applied the PCA and K-means model to the test set:

- All cells and microenvironments from test set samples are extracted;
- SPACE-GM is applied to test set microenvironments to extract embeddings;
- PCA and K-means models trained with the reference dataset are directly applied to test set microenvironment embeddings, their cluster assignments are collected and summarized.

For PCA and K-means, we used the Python implementations from `scikit-learn`^44^: `sklearn.decomposition.PCA` and `sklearn.cluster.KMeans`. For UMAP, we used the Python implementation from `umap-learn`^48^.

### In silico permutation of cells in microenvironments

Analysis of SPACE-GM embeddings uncovered groups of microenvironments that are disease-relevant; we then applied permutation experiments on microenvironments-of-interest to discover and validate structural motifs that are indicative of phenotypes.

Two general forms of permutations are implemented:

- Dispersed permutation:
  ○ A list of target cell types are provided;
  ○ On the microenvironment/patch/sample (jointly denoted as cellular graphs), identify all cells whose cell type appears in the target list and record their spatial location and other cellular features;
  ○ Randomly permute the list of cells identified in the previous step, then assign the permuted cell types and cellular features back to the original cellular graph;
  ○ Run (trained) SPACE-GM on the permuted cellular graph and perform microenvironment-aggregation if needed;
- Coherent permutation:
  ○ A list of target cell types and the center coordinate of cellular graph are provided;
  ○ On the cellular graph, identify all cells whose cell type appears in the target list and record their spatial location and other cellular features;
  ○ Sort the list of cells identified in the previous step by cell type;
  ○ Calculate polar coordinates for the list of cells using the cellular graph center as the pole;
  ○ Assign cell types and cellular features back to the original cellular graph sequentially following the order of azimuthal angle. Resulting permuted cellular graphs should have cells of the same type appearing in the same sector around the graph center.

In the experiment on heterogeneous tumor microenvironments, we performed permutations on patches and whole samples. Patches are selected following the procedure below:

- On each of the UPMC-HNC samples, we derive the cluster assignment of every individual cell following the dimensionality reduction and clustering pipeline introduced above;
- We iterate through the list of all cells that are assigned to the heterogeneous tumor microenvironment until we find a cell that satisfies the following criterion. Note that up to one patch can be selected per sample.
  ○ Isolate all cells that are within a distance of 185 μm to the query cell (denoted as a patch);
  ○ More than 40% of the cells in the patch are assigned to heterogeneous tumor microenvironments;
- Extract the query cell and its surrounding patch. For all cells in the patch that are within a distance of 110 μm to the center (to guarantee the completeness of the 3-hop neighborhood), extract their microenvironments and perform predictions.

61 patches are selected, from which we picked the 50 patches that have higher entropy (of cell type frequency vector). Permutations are performed on the following cell types: *Tumor 2*, *Tumor 4*, *Tumor 5 (Ki67+)*, and *Tumor 6*, in which the interaction between *Tumor 4* and *Tumor 5 (Ki67+)* has the biggest influence.

In the experiment on granulocyte-tumor microenvironments, we performed coherent and dispersed permutations on different microenvironment groups to reverse the organization pattern of granulocytes. We didn’t perform patch-level permutation due to difficulty in finding regional patches rich in either microenvironment. All cell types are included in the target list and permuted in this experiment. Note that in the coherent permutation case, all cells except for tumor cells (*Tumor 1* through *Tumor 6*) are aligned coherently according to the procedure above, tumor cells are first combined and randomly permuted before aligning to avoid confounding. See Supplementary Fig. 7 for example.

## Acknowledgements

J.Z. is supported by NSF CAREER 1942926. K.S. is supported by a Knight-Hennessy Fellowship.

## Data and Code Availability

Codes are available at https://gitlab.com/enable-medicine-public/space-gm. Data (CODEX images and graphical representations) will be available upon request.

## Supplementary Tables and Figures

**Supplementary Table 1.**
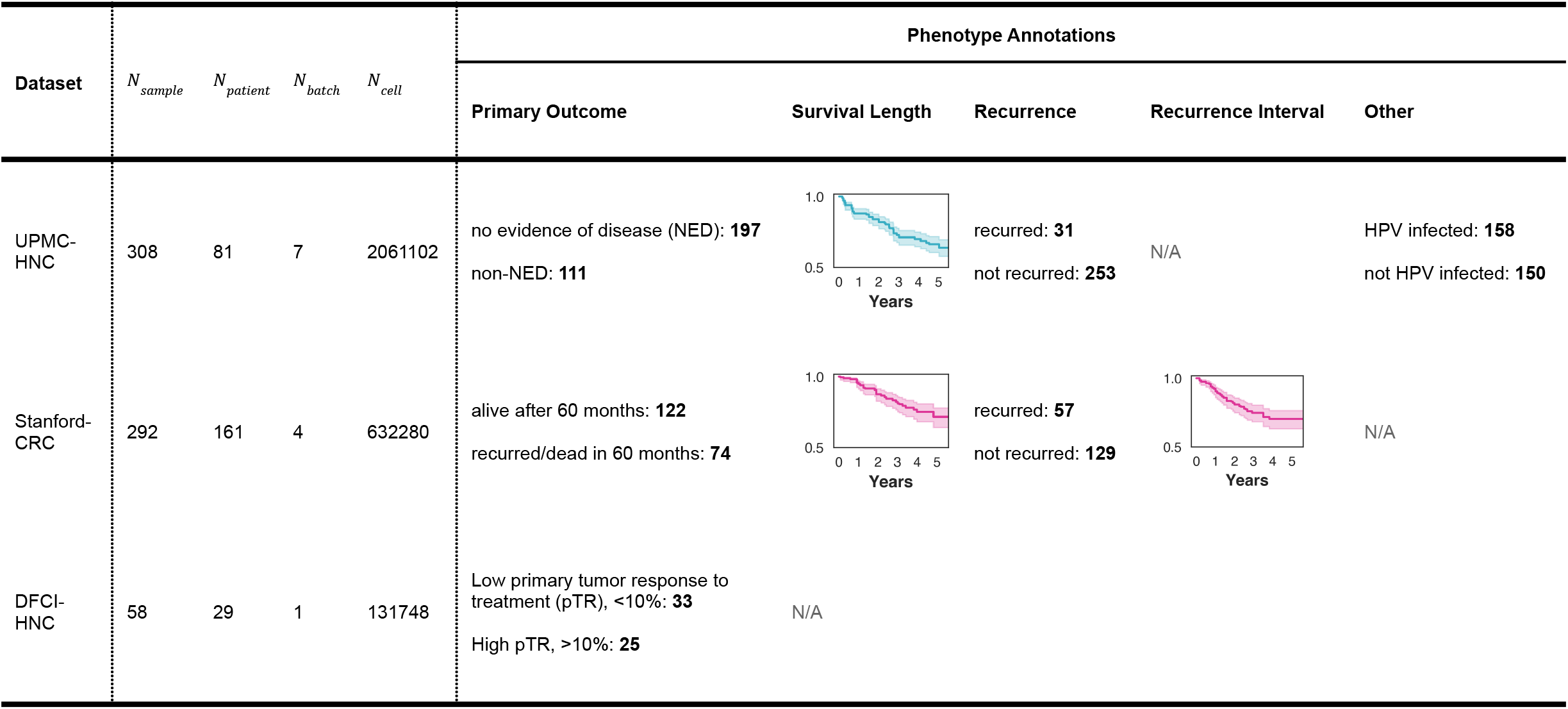
Details of datasets and phenotype annotations. Tasks “Primary Outcome”, “Recurrence” and “Other” have binary labels and are used for binary classification. Tasks “Survival Length” and “Recurrence Interval” provide time-to-event data and are used for hazards modeling.

| Dataset | $N_{sample}$ | $N_{patient}$ | $N_{batch}$ | $N_{cell}$ | Phenotype Annotations | | | | |
| --- | --- | --- | --- | --- | --- | --- | --- | --- | --- |
|  |  |  |  |  | Primary Outcome | Survival Length | Recurrence | Recurrence Interval | Other |
| UPMC-HNC | 308 | 81 | 7 | 2061102 | no evidence of disease (NED): **197**<br>non-NED: **111** |  | recurred: **31**<br>not recurred: **253** | N/A | HPV infected: **158**<br>not HPV infected: **150** |
| Stanford-CRC | 292 | 161 | 4 | 632280 | alive after 60 months: **122**<br>recurred/dead in 60 months: **74** |  | recurred: **57**<br>not recurred: **129** |  | N/A |
| DFCI-HNC | 58 | 29 | 1 | 131748 | Low primary tumor response to treatment (pTR), <10%: **33**<br>High pTR, >10%: **25** | N/A |  |  |  |

**Supplementary Table 2.** Prediction performances on UPMC tasks: four-fold patient cross-validation, average performances over all four folds are reported.

| Model | Binary Classification (ROC-AUC) |  |  | Hazards Model (C-index) |
| --- | --- | --- | --- | --- |
|  | HPV Infection | Primary Outcome | Recurrence | Survival Length |
| Linear on composition (sample) | 0.830 | 0.739 | 0.702 | 0.694 |
| MLP on composition (sample) | 0.757 | 0.711 | 0.672 | 0.669 |
| Linear on composition (microenvironment) | 0.852 | 0.749 | 0.720 | 0.700 |
| MLP on composition (microenvironment) | 0.863 | 0.743 | 0.706 | 0.693 |
| SPACE-GM no-pretraining | 0.895 | 0.756 | 0.755 | 0.705 |
| SPACE-GM | 0.900 | 0.760 | 0.766 | 0.722 |

**Supplementary Table 3.** Prediction uncertainty on UPMC tasks: on each of the two cross-coverslip validation folds, we randomly select validation samples (with replacement) and evaluate model performances. Means and standard deviations of 10 independently bootstrapped runs are reported.

| Model | Validation Fold | Binary Classification (ROC-AUC) |  |  | Hazards Model (C-index) |
| --- | --- | --- | --- | --- | --- |
|  |  | HPV Infection | Primary Outcome | Recurrence | Survival Length |
| Linear on composition (sample) | 0 | 0.850 ± 0.030 | 0.773 ± 0.035 | 0.840 ± 0.045 | 0.700 ± 0.032 |
|  | 1 | 0.873 ± 0.022 | 0.768 ± 0.031 | 0.826 ± 0.040 | 0.684 ± 0.026 |
| MLP on composition (sample) | 0 | 0.851 ± 0.029 | 0.789 ± 0.033 | 0.888 ± 0.040 | 0.694 ± 0.053 |
|  | 1 | 0.848 ± 0.033 | 0.760 ± 0.028 | 0.794 ± 0.040 | 0.704 ± 0.030 |
| Linear on composition (microenvironment) | 0 | 0.854 ± 0.029 | 0.775 ± 0.041 | 0.816 ± 0.053 | 0.706 ± 0.057 |
|  | 1 | 0.876 ± 0.032 | 0.775 ± 0.043 | 0.818 ± 0.066 | 0.695 ± 0.042 |
| MLP on composition (microenvironment) | 0 | 0.885 ± 0.026 | 0.841 ± 0.048 | 0.867 ± 0.038 | 0.768 ± 0.054 |
|  | 1 | 0.903 ± 0.023 | 0.790 ± 0.048 | 0.827 ± 0.043 | 0.734 ± 0.021 |
| SPACE-GM no-pretraining | 0 | 0.912 ± 0.017 | 0.885 ± 0.025 | 0.906 ± 0.057 | 0.798 ± 0.033 |
|  | 1 | 0.917 ± 0.018 | 0.835 ± 0.019 | 0.874 ± 0.033 | 0.766 ± 0.027 |
| SPACE-GM | 0 | 0.928 ± 0.020 | 0.902 ± 0.020 | 0.921 ± 0.048 | 0.814 ± 0.027 |
|  | 1 | 0.928 ± 0.017 | 0.845 ± 0.022 | 0.876 ± 0.028 | 0.788 ± 0.022 |

**Supplementary Figure 1.**
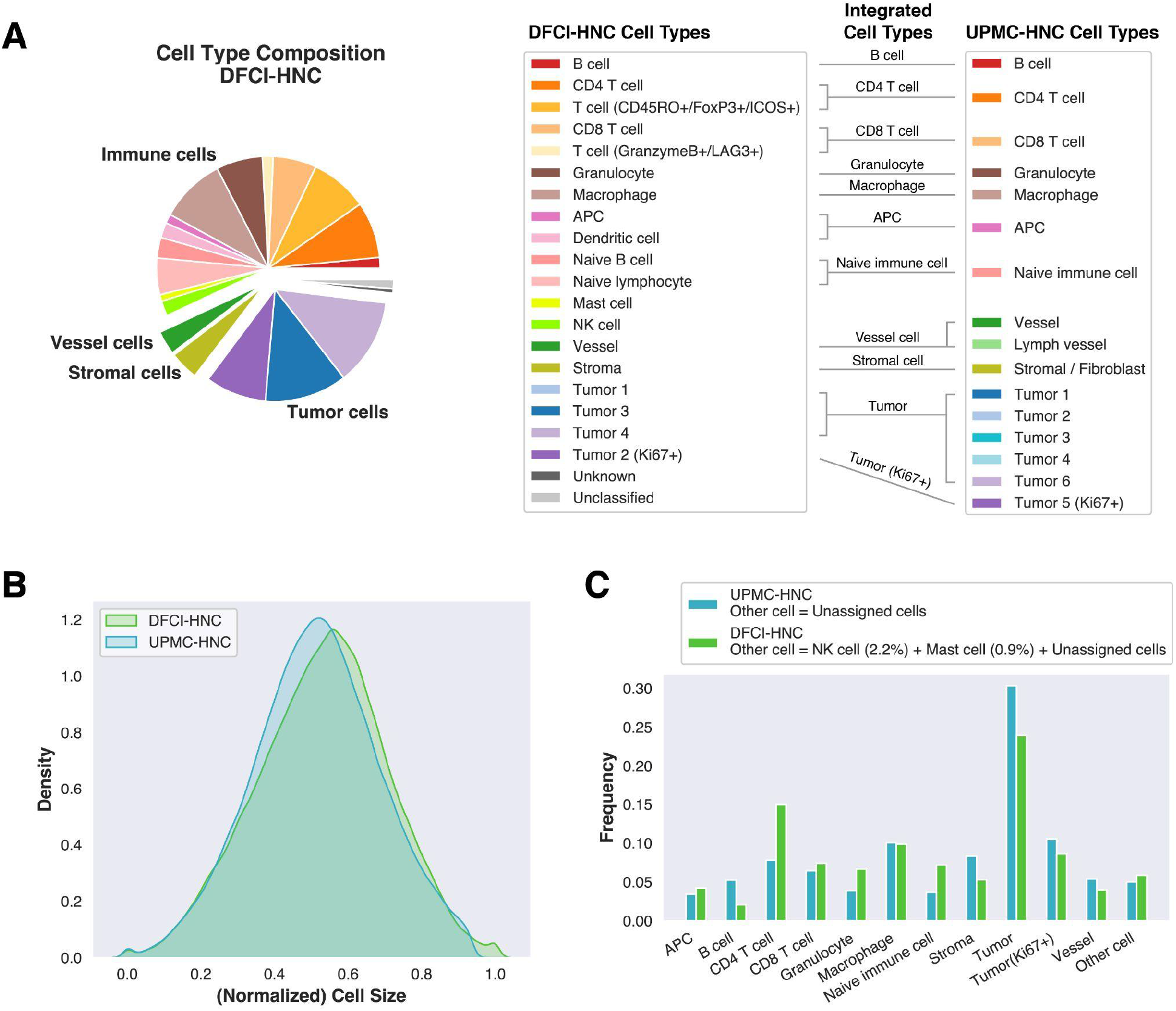
DFCI-HNC cell characteristics. **(A)** Cell type composition of the DFCI-HNC dataset. We further match the cell type to the UPMC-HNC dataset. **(B)** Distribution of normalized cell sizes (Methods) in UPMC-HNC and DFCI-HNC samples. **(C)** Composition of integrated cell types in UPMC-HNC and DFCI-HNC samples. Note that all tumor subtypes (except for Ki67+ subtype) are combined into the “Tumor” category; certain cell types (NK cell, mast cell) that are unique to one of the datasets are moved to the “Other cell” category.

**Supplementary Figure 2.**
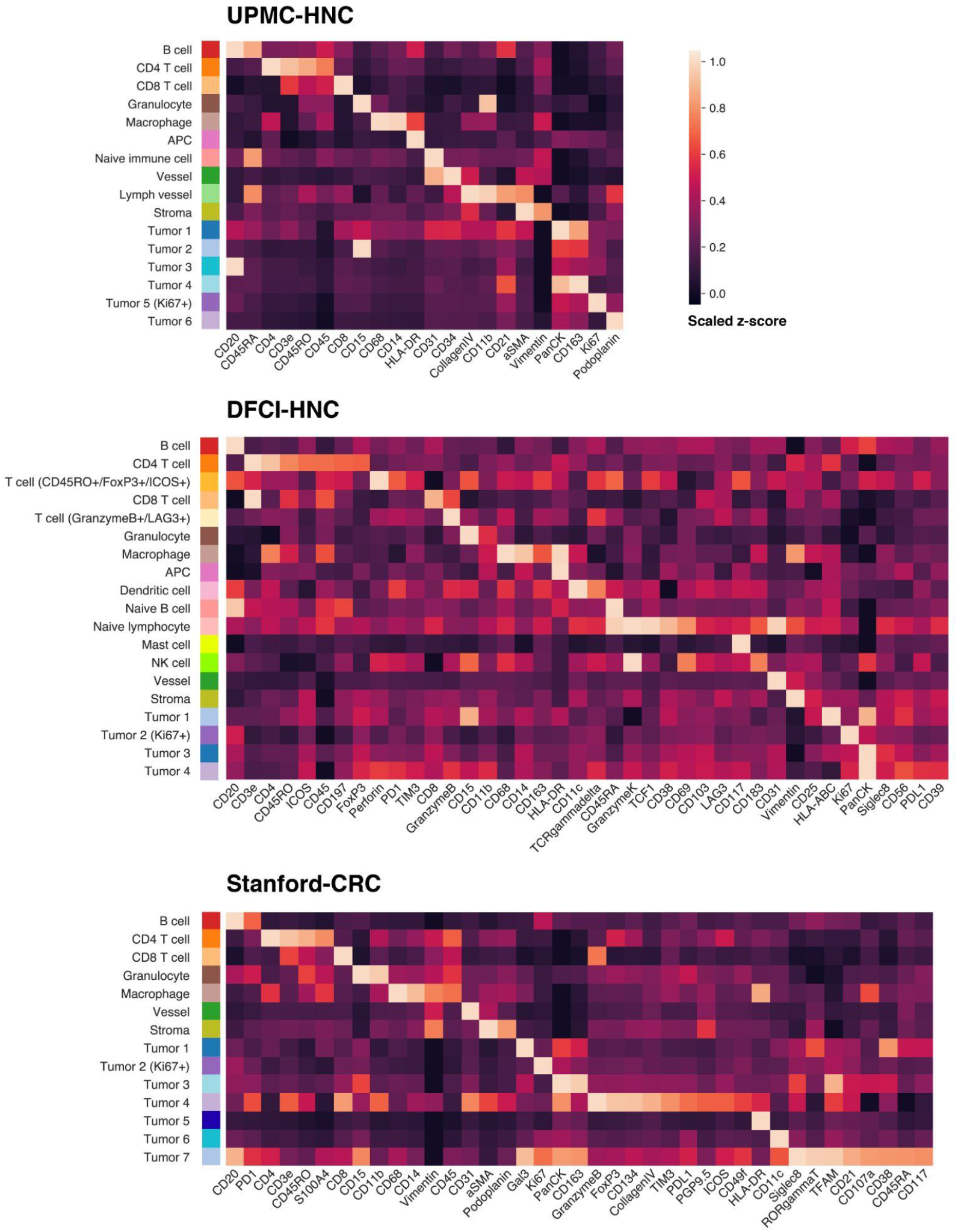
Marker expression of different cell types. Heatmaps of average marker expression for each cell type in UPMC-HNC (top row), DFCI-HNC (middle row) and Stanford-CRC (bottom row) are visualized. Note that each dataset uses a different marker panel.

**Supplementary Figure 3.**
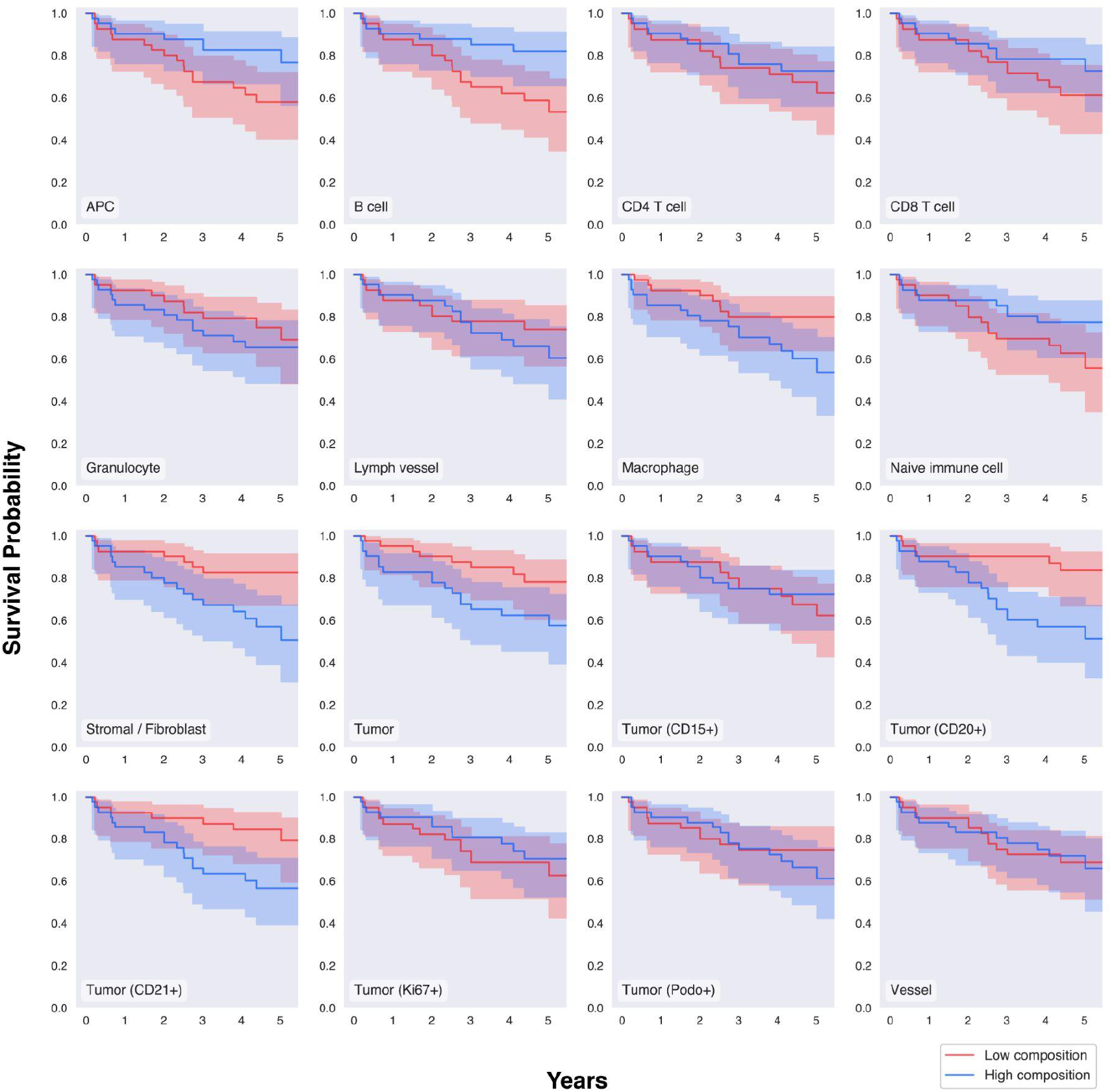
Survival curves of different cell type composition groups. In each of the 16 panels, we separate all test samples into two groups based on the composition of the corresponding cell type. The survival curves of the two groups are visualized. Note that the separation between groups is much weaker than the risk-based grouping in Figure 3C.

**Supplementary Figure 4.**
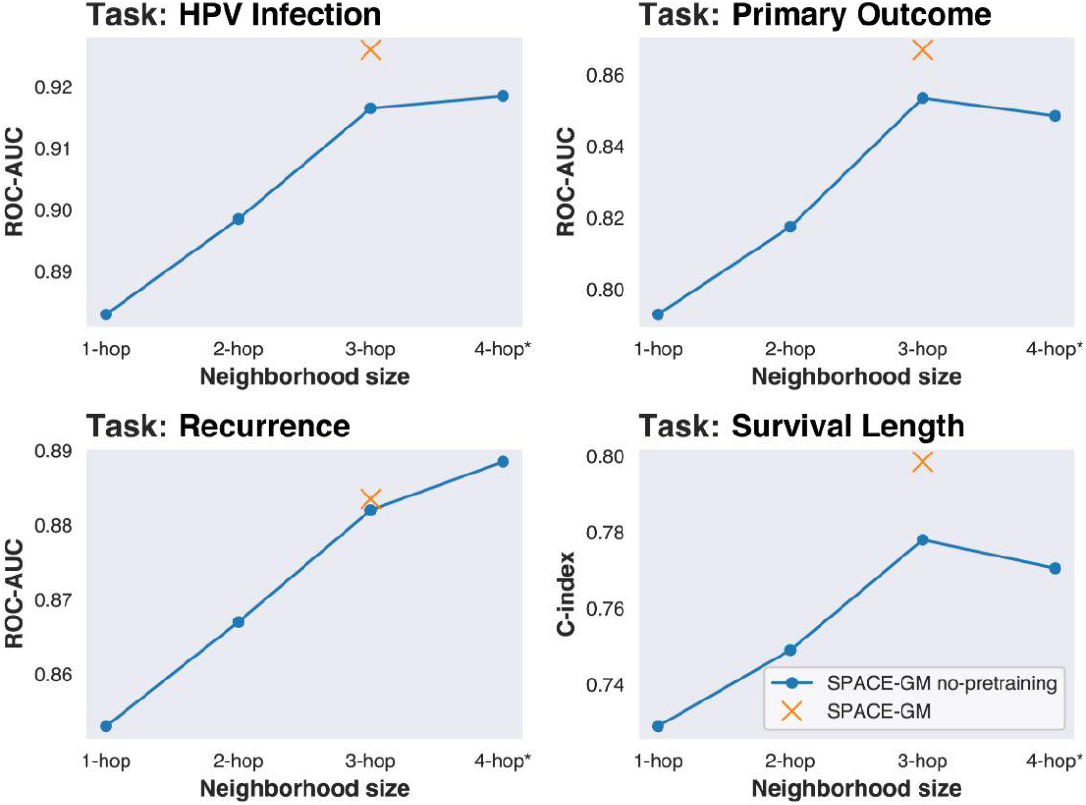
Performance on microenvironments of different sizes. We trained a series of SPACE-GM no-pretraining models with microenvironment inputs of different sizes (from 1-hop to 4-hop). Three-layer GINs are used in cases of 1-hop, 2-hop, and 3-hop inputs. Four-layer GIN is used for 4-hop inputs. Results show that performances increase monotonically before 3-hop and plateau around 3-hop and 4-hop.

**Supplementary Figure 5.**
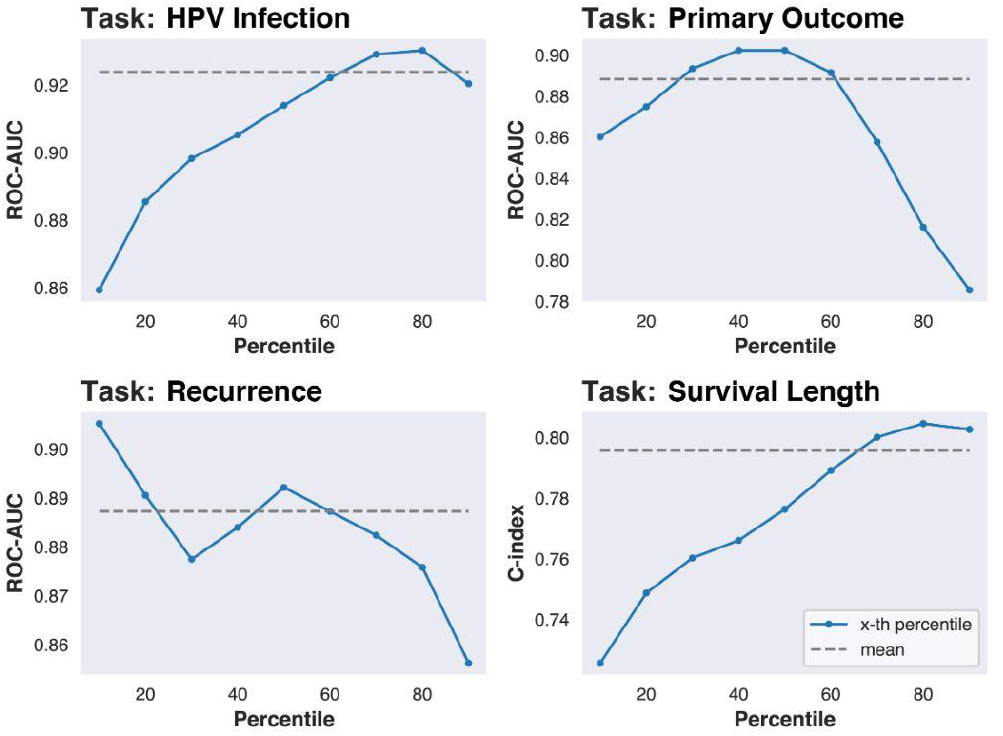
Performance of different aggregation thresholds. Microenvironment-aggregation is a necessary step to calculate whole-sample predictions for SPACE-GM (and other microenvironment-based methods). On one of the validation folds of UPMC-HNC, we evaluate performances on predictions of different percentile thresholds. Dashed lines mark the performance of mean-aggregated predictions. We conclude that mean aggregation provides robust performances.

**Supplementary Figure 6.**
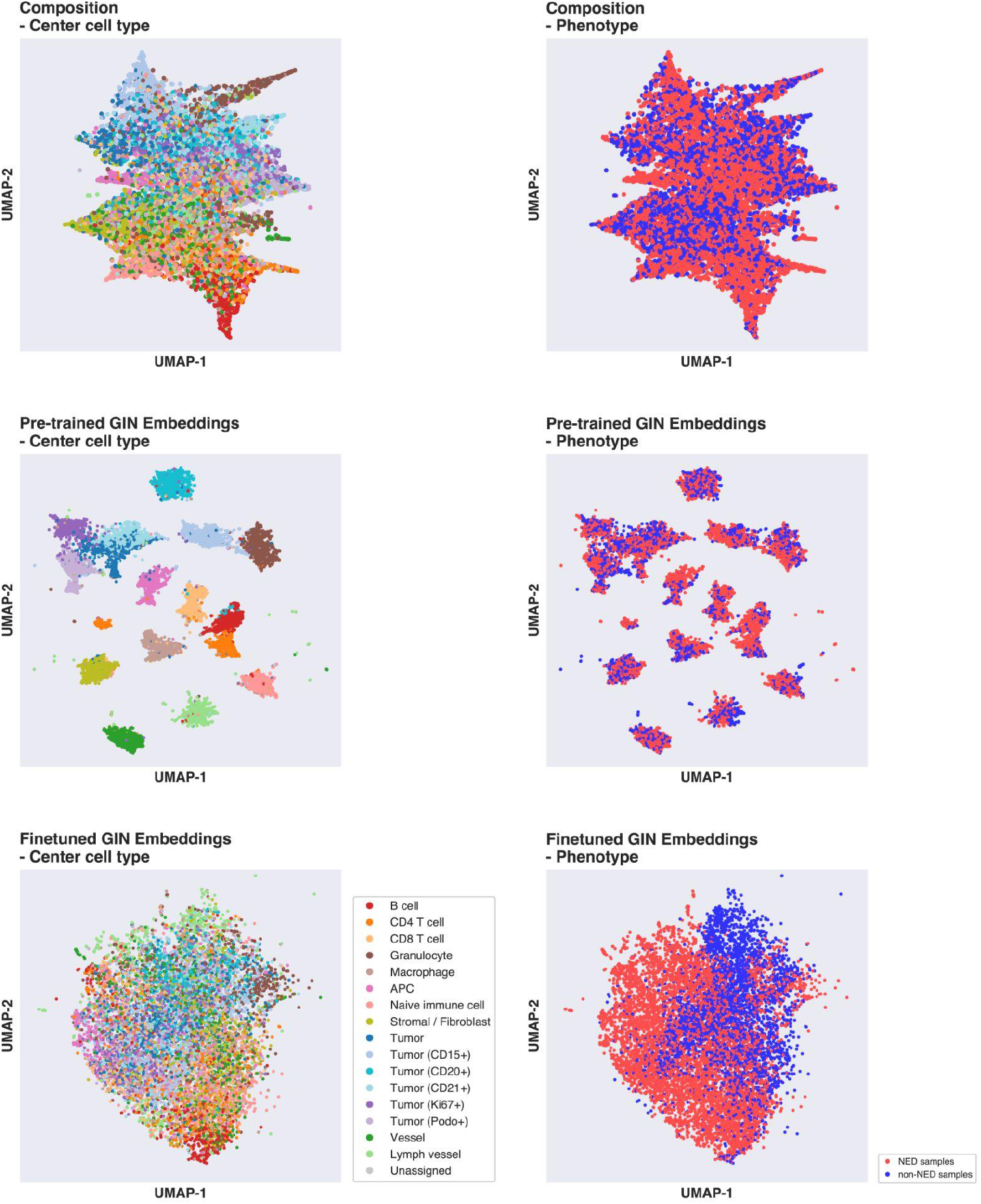
UMAP of composition vectors and microenvironment embeddings. Embeddings are extracted after each of the two training stages of SPACE-GM (pre-training on cellular expression and finetuning on primary outcome task). Embeddings from the pre-trained model show strong correlations with center cell type. Note that the clustering captured the similarity between cell types: lymphocytes appear together; tumor cells form a large cluster. Embeddings from the final finetuned SPACE-GM model correlate more with phenotype, as we see a clear blue-red color separation in the distribution.

**Supplementary Figure 7.**
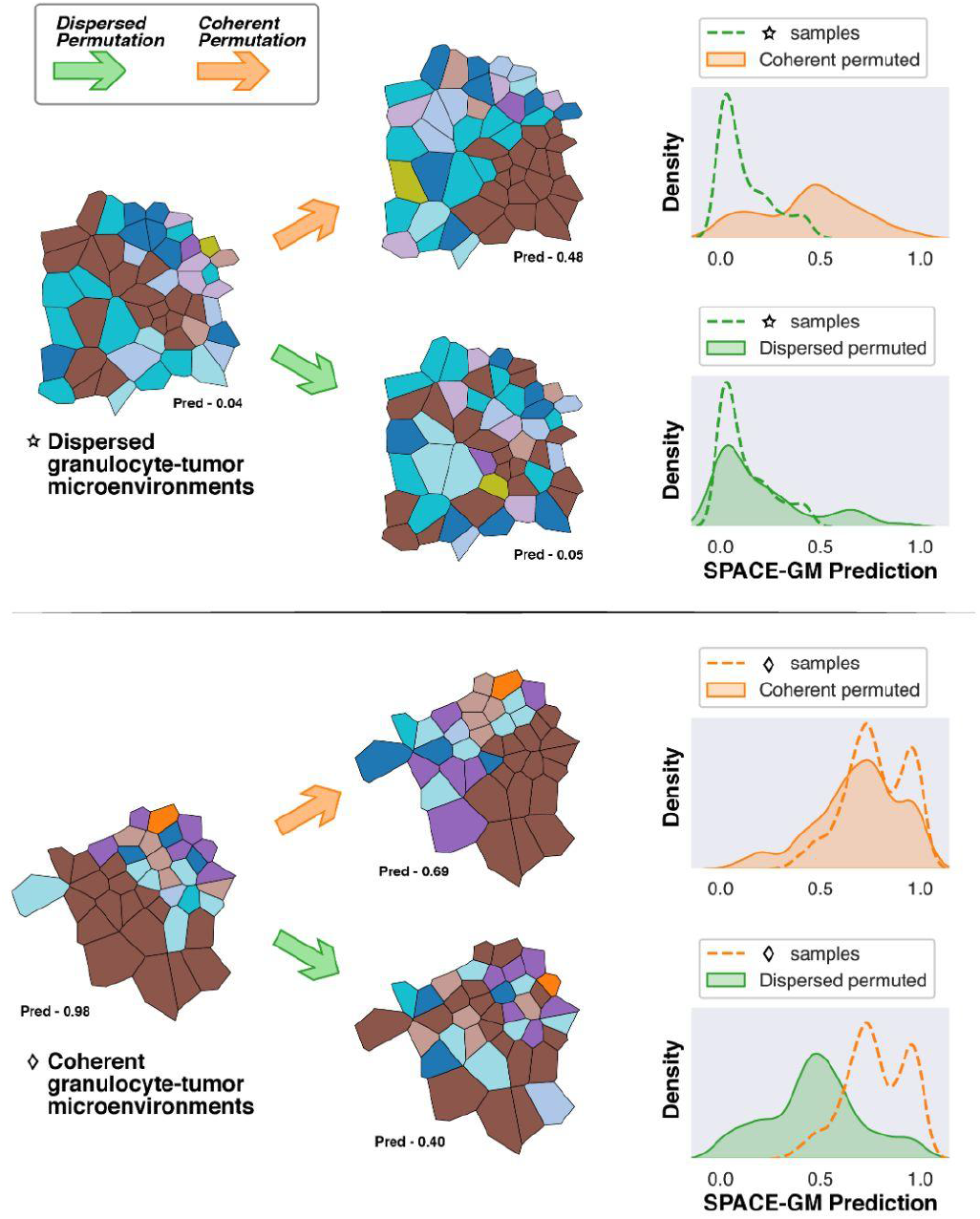
In silico permutation of granulocyte-tumor microenvironments. On the two granulocyte-tumor rich microenvironments, we perform coherent and dispersed permutations that could change the local arrangement pattern of granulocytes (Methods). We observe significant shifts in the distribution of SPACE-GM predictions when applying coherent permutation on the dispersed group (and vice versa), indicating that the dispersion of granulocytes affects model predictions and can have potential correlations with patient outcomes.

**Supplementary Figure 8.**
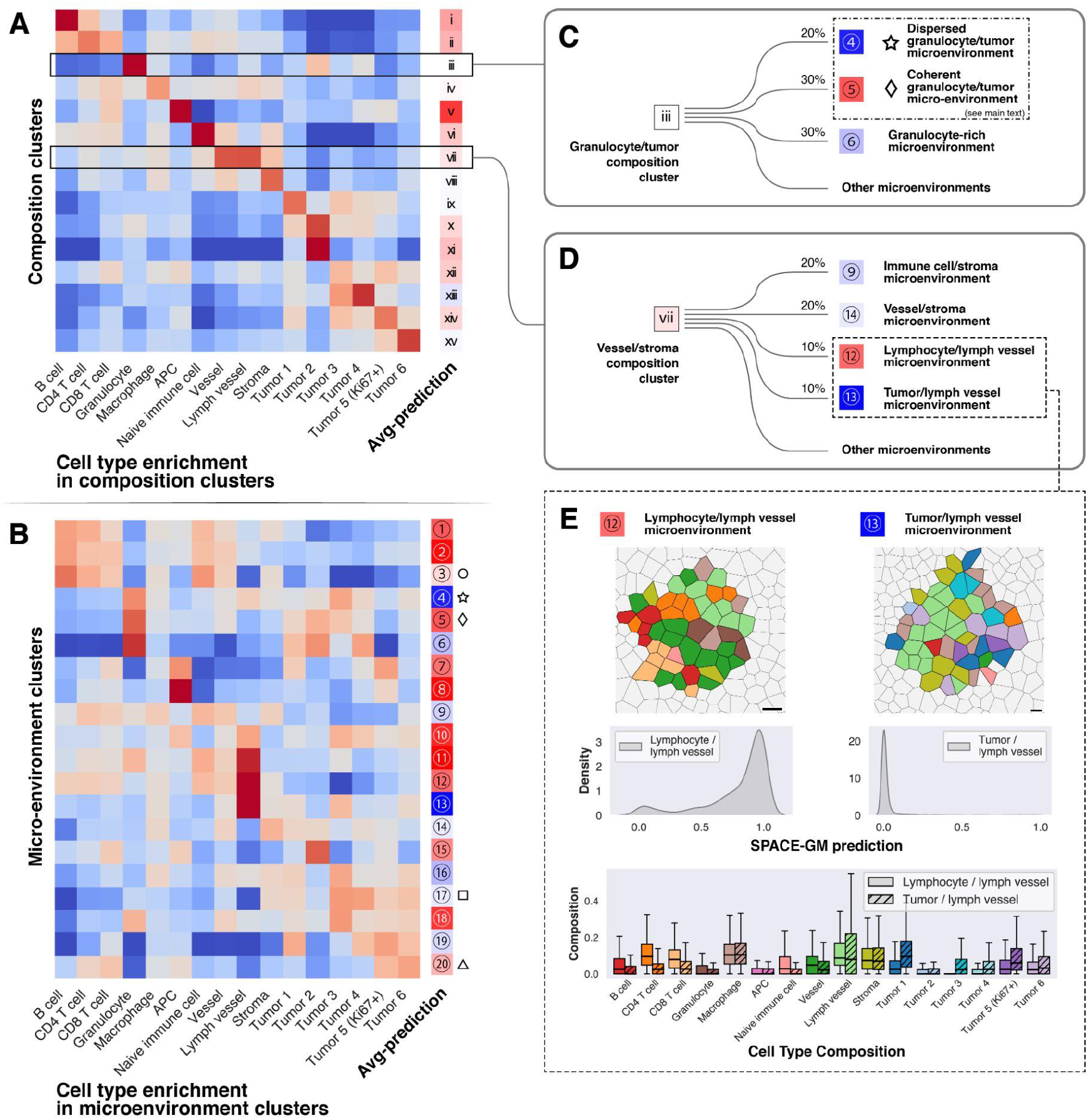
Comparison of composition-based clusters and microenvironment clusters. **(A)** and **(B)** We generate composition-based clusters with cell type compositions of 3-hop subgraphs (microenvironments) following the same procedure. Note that the average predictions of microenvironment clusters are much more polarized. **(C)** The composition cluster highly enriched with granulocytes has neutral average predictions/labels, it could be further dissected into multiple sub-clusters that belong to different microenvironment clusters. We see a pair of granulocyte/tumor microenvironments that have opposite outcome labels, see (Results) for further discussion. **(D)** Similarly, the composition cluster enriched with vessel/lymph vessel cells could be dissected into multiple sub-clusters, from which we notice two microenvironment clusters that are both enriched with lymph vessel cells but have different composition and outcome predictions. **(E)** Comparison of the two microenvironments enriched in lymph vessel cells: the left column shows a microenvironment with more lymphocytes and has overall positive outcomes; the right column shows a contrasting group with more tumor cells and much worse outcome predictions. Observation of tumor cells in close vicinity of lymph vessels indicates potential lymphovascular invasion and will lead to worse prognosis, which aligns with model predictions.

## Supplementary Notes

### Microenvironments of different sizes

SPACE-GM constructs microenvironments based on n-hop neighborhoods. To balance prediction performance and microenvironment size (which is directly related to ease of interpretability), we benchmark microenvironments of different sizes in their prediction performances.

We define n-hop neighborhoods as all the nodes reachable from the query nodes within n edges: 1-hop neighborhoods only contain immediate neighbors, and 2-hop neighborhoods contain neighboring cells of the immediate neighbors. In this experiment, we train a series of SPACE-GM models with different sizes of inputs by enumerating neighborhood radius (n). For ease of computation, we only compare performances of SPACE-GM no-pretraining models. Supplementary Fig. 4 illustrates the phenotype prediction performance with regards to input microenvironment size, in which the same 3-layer GIN model structure is used for 1-hop, 2-hop, and 3-hop inputs, while a 4-layer GIN model is used for 4-hop inputs.

We notice that performances increase monotonically before 3-hop, suggesting that a larger microenvironment is indeed beneficial for modeling phenotypic properties. 4-hop models are slightly better on two out of the four tasks evaluated, but do not present a strong overall advantage. Given that microenvironment size grows quadratically with increasing neighborhood radius, using inputs larger than 3-hop might cause difficulties in the following structural and permutation analysis, so we choose to work with microenvironments defined as 3-hop neighborhoods.

### Microenvironment-aggregation methods

SPACE-GM is trained on subgraphs/microenvironments. To evaluate final prediction performance, we need a *post hoc* procedure that aggregates predictions of all microenvironments from a single sample. For this step, we evaluate aggregation functions including mean, median, and a series of quantile functions at different thresholds (Supplementary Fig. 5). Plots show that different tasks have different preferences for quantile thresholds; survival length has the best performance when using the 80-th percentile, indicating that within each slice nodes with high risk predictions are more correlated with actual survival probabilities. While there are no clear clues on a universally suitable threshold for quantiles, averaging microenvironment predictions is usually as good as the best quantile thresholds. Therefore, mean-aggregation is used as the default method for microenvironment-aggregation.

### Cross-study application of SPACE-GM

We test the generalizability of SPACE-GM by applying models trained with UPMC-HNC samples to DFCI-HNC. In this procedure, due to the independent preprocessing pipelines for both datasets, we need to integrate cellular features (node features) before deploying the model.

SPACE-GM uses cellular features including cell type and cell size (Methods). While cell size is calculated based on the size of segmentation masks, cell type is derived through a per-dataset unsupervised PCA-kNN procedure (Methods). Supplementary Fig. 1A illustrates the cell classification results for DFCI-HNC, which shared many common cell type populations with UPMC-HNC; we further define integrated cell types between the two datasets to unify input features to SPACE-GM. Note that most tumor subpopulations are combined to avoid confusion, except that *Tumor (Ki67+)* is separated and treated as the proliferating tumor sub-population. It is worth noting that integrating cell types hurts SPACE-GM prediction performance due to the decrease in the heterogeneity of inputs.

After integration, we compare the input features of UPMC-HNC and DFCI-HNC datasets. Supplementary Fig. 1B compares the distribution of normalized cell size (Methods), in which the difference is minimal. Supplementary Fig. 1C compares the distribution of integrated cell types, DFCI-HNC has more CD4 T cells and fewer B cells, but the overall balance between immune cells and tumor cells is similar to UPMC-HNC. Note that we assign minor cell populations (e.g., Mast cell) not present in both datasets to the “Other cell” group.

### Benchmark prediction uncertainty

The main evaluation experiment of SPACE-GM compares its performance against baseline composition-based methods on a series of patient phenotype prediction tasks using two independent cross-coverslip validation folds. To study the uncertainty of model prediction performances, we calculate the standard deviations (SDs) of performance metrics by bootstrapping.

In Supplementary Table 3, we evaluate model prediction performances by bootstrapping validation samples (random resampling with replacement). 10 independent runs are conducted, from which means and SDs are calculated and reported. In most test cases, SPACE-GM performance exceeds the best-performing composition-based model by at least 2 SD, indicating a robust advantage over baseline methods.

### Parallel composition-based clusters and microenvironment clusters

In the main text, we discussed clustering using microenvironment embeddings derived from SPACE-GM models trained with phenotypic property prediction. The resulting clusters are intrinsically correlated with target phenotypes. To further validate this argument, we perform the same clustering analysis on composition inputs, excluding any phenotype-guided supervision.

We apply the same dimensionality reduction and clustering procedure as introduced in (Methods) on composition vectors of the same set of reference microenvironments. Instead of using the top 20 PCs, we used the top PCs (12) that capture 90% of the total variance in compositions. K-means clustering is adjusted accordingly (K = 15).

Supplementary Fig. 8A plots the enrichment of cell types and average SPACE-GM predictions (very similar to labels) for each cluster. Compared with Supplementary Fig. 8B (microenvironment cluster characteristics), it is obvious that composition-based clusters are more coherent in terms of cell types: most of the clusters are highly enriched for one or two cell types while having fewer other cells. On the other hand, average predictions of microenvironment clusters are more polarized than composition-based ones, facilitating the discovery of disease-relevant clusters.

We further look at two concrete examples: Supplementary Fig. 8C shows a granulocyte-rich composition cluster that has overall neutral average predictions or labels. If we look at the microenvironment cluster assignments, this composition-based cluster can be dissected into multiple subgroups belonging to different microenvironment clusters. More specifically, we notice that it maps to two microenvironment clusters ④ and ⑤, which are exactly the pair we discussed with permutation experiments. They have similar composition but opposite phenotypic predictions, which can be partially attributed to the dispersion of granulocytes.

#### Microenvironments enriched in lymph vessel cells

In the other example, we look at the composition-based cluster enriched in *Vessel* and *Lymph vessel* cells (Supplementary Fig. 8D). We similarly parallel it with microenvironment clusters and found a pair of contrasting groups that are both enriched in *lymph vessel* cells. They display different cell type compositions other than *Lymph vessel* (Supplementary Fig. 8E): cluster ⑫ has more immune cells including *B cell*, *CD4 T cell*, etc., while cluster ⑬ is more enriched in *Tumor 1*, *Tumor 5 (Ki67+)*. We hypothesize that cluster ⑫ represents normal lymph vessel structure, while cluster ⑬ might indicate lymphovascular invasion, a known negative factor for prognosis.

Though such differences can already be captured with composition vectors, their distinction will likely be overwhelmed by major variance such as the amount of dominating cell population (i.e., *Lymph vessel* cells) when applying unsupervised clustering. However, when clustering on phenotype-guided embeddings, key distinctions captured by the neural network will be enlarged in the embedding space and hence more easily perceived in the downstream analysis.

## Reference

1. Atlas of clinically distinct cell states and ecosystems across human solid tumors. Cell 184, 5482–5496.e28 (2021).

2. Wu, S. Z. et al. A single-cell and spatially resolved atlas of human breast cancers. Nat. Genet. 53, 1334–1347 (2021).

3. Bejarano, L., Jordāo, M. J. C. & Joyce, J. A. Therapeutic Targeting of the Tumor MicroenvironmentTherapeutic Targeting of the Tumor Microenvironment. Cancer Discov. 11, 933–959 (2021).

4. Lomakin, A. et al. Spatial genomics maps the structure, character and evolution of cancer clones. doi:10.1101/2021.04.16.439912.

5. Rodriques, S. G. et al. Slide-seq: A scalable technology for measuring genome-wide expression at high spatial resolution. Science 363, 1463–1467 (2019).

6. Wang, X. et al. Three-dimensional intact-tissue sequencing of single-cell transcriptional states. Science 361, (2018).

7. Moffitt, J. R. et al. High-throughput single-cell gene-expression profiling with multiplexed error-robust fluorescence in situ hybridization. Proceedings of the National Academy of Sciences vol. 113 11046–11051 (2016).

8. Goltsev, Y. et al. Deep Profiling of Mouse Splenic Architecture with CODEX Multiplexed Imaging. Cell 174, 968–981.e15 (2018).

9. Angelo, M. et al. Multiplexed ion beam imaging of human breast tumors. Nat. Med. 20, 436–442 (2014).

10. Lin, J.-R., Fallahi-Sichani, M., Chen, J.-Y. & Sorger, P. K. Cyclic Immunofluorescence (CycIF), A Highly Multiplexed Method for Single-cell Imaging. Curr. Protoc. Chem. Biol. 8, 251–264 (2016).

11. Ali, H. R. et al. Imaging mass cytometry and multiplatform genomics define the phenogenomic landscape of breast cancer. Nat Cancer 1, 163–175 (2020).

12. Schürch, C. M. et al. Coordinated Cellular Neighborhoods Orchestrate Antitumoral Immunity at the Colorectal Cancer Invasive Front. Cell 183, 838 (2020).

13. Bhate, S. S., Barlow, G. L., Schürch, C. M. & Nolan, G. P. Tissue schematics map the specialization of immune tissue motifs and their appropriation by tumors. Cell Syst 13, 109–130.e6 (2022).

14. Zhou, Y. et al. Cgc-net: Cell graph convolutional network for grading of colorectal cancer histology images. in Proceedings of the IEEE/CVF International Conference on Computer Vision Workshops 0–0 (2019).

15. Lu, W., Graham, S., Bilal, M., Rajpoot, N. & Minhas, F. Capturing cellular topology in multi-gigapixel pathology images. in Proceedings of the IEEE/CVF Conference on Computer Vision and Pattern Recognition Workshops 260–261 (2020).

16. Anand, D., Gadiya, S. & Sethi, A. Histographs: graphs in histopathology. in Medical Imaging 2020: Digital Pathology vol. 11320 150–155 (SPIE, 2020).

17. Gilmer, J., Schoenholz, S. S., Riley, P. F., Vinyals, O. & Dahl, G. E. Neural Message Passing for Quantum Chemistry. in Proceedings of the 34th International Conference on Machine Learning (eds. Precup, D. & Teh, Y. W.) vol. 70 1263–1272 (PMLR, 06--11 Aug 2017).

18. Wu, Z. et al. A Comprehensive Survey on Graph Neural Networks. IEEE Trans Neural Netw Learn Syst 32, 4–24 (2021).

19. Brbić, M. et al. Annotation of Spatially Resolved Single-cell Data with STELLAR. doi:10.1101/2021.11.24.469947.

20. Innocenti, C. et al. An Unsupervised Graph Embeddings Approach to Multiplex Immunofluorescence Image Exploration. doi:10.1101/2021.06.09.447654.

21. Fischer, D. S., Schaar, A. C. & Theis, F. J. Learning cell communication from spatial graphs of cells. doi:10.1101/2021.07.11.451750.

22. Kim, J. et al. Unsupervised discovery of tissue architecture in multiplexed imaging. doi:10.1101/2022.03.15.484534.

23. Xu, K., Hu, W., Leskovec, J. & Jegelka, S. How Powerful are Graph Neural Networks? arXiv [cs.LG] (2018).

24. Argiris, A., Karamouzis, M. V., Raben, D. & Ferris, R. L. Head and neck cancer. Lancet 371, 1695–1709 (2008).

25. Dalerba, P. et al. CDX2 as a Prognostic Biomarker in Stage II and Stage III Colon Cancer. N. Engl. J. Med. 374, 211–222 (2016).

26. Uppaluri, R. et al. Neoadjuvant and Adjuvant Pembrolizumab in Resectable Locally Advanced, Human Papillomavirus–Unrelated Head and Neck Cancer: A Multicenter, Phase II Trial. Clinical Cancer Research vol. 26 5140–5152 (2020).

27. Blise, K. E., Sivagnanam, S., Banik, G. L., Coussens, L. M. & Goecks, J. Single-cell spatial architectures associated with clinical outcome in head and neck squamous cell carcinoma. NPJ Precis Oncol 6, 10 (2022).

28. Jackson, H. W. et al. The single-cell pathology landscape of breast cancer. Nature 578, 615–620 (2020).

29. Trellakis, S. et al. Polymorphonuclear granulocytes in human head and neck cancer: enhanced inflammatory activity, modulation by cancer cells and expansion in advanced disease. Int. J. Cancer 129, 2183–2193 (2011).

30. Lonardi, S. et al. Tumor-associated neutrophils (TANs) in human carcinoma-draining lymph nodes: a novel TAN compartment. Clin Transl Immunology 10, e1252 (2021).

31. Coffelt, S. B. Wellenstein, M. D. & de Visser, K. E. Neutrophils in cancer: neutral no more. Nat. Rev. Cancer 16, 431–446 (2016).

32. Shojaei, F. et al. Bv8 regulates myeloid-cell-dependent tumour angiogenesis. Nature 450, 825–831 (2007).

33. Di Mitri, D. et al. Tumour-infiltrating Gr-1+ myeloid cells antagonize senescence in cancer. Nature 515, 134–137 (2014).

34. Coffelt, S. B. et al. IL-17-producing γδ T cells and neutrophils conspire to promote breast cancer metastasis. Nature 522, 345–348 (2015).

35. Wculek, S. K. & Malanchi, I. Neutrophils support lung colonization of metastasis-initiating breast cancer cells. Nature 528, 413–417 (2015).

36. Regev, A. et al. The Human Cell Atlas. Elife 6, (2017).

37. HuBMAP Consortium. The human body at cellular resolution: the NIH Human Biomolecular Atlas Program. Nature 574, 187–192 (2019).

38. Greenwald, N. F. et al. Whole-cell segmentation of tissue images with human-level performance using large-scale data annotation and deep learning. Nat. Biotechnol. 40, 555–565 (2021).

39. Hickey, J. W., Tan, Y., Nolan, G. P. & Goltsev, Y. Strategies for Accurate Cell Type Identification in CODEX Multiplexed Imaging Data. Front. Immunol. 12, 727626 (2021).

40. Blondel, V. D., Guillaume, J.-L., Lambiotte, R. & Lefebvre, E. Fast unfolding of communities in large networks. (2008) doi:10.1088/1742-5468/2008/10/P10008.

41. Geovoronoi. PyPI https://pypi.org/project/geovoronoi/.

42. Kvamme, H., Borgan, Ø. & Scheel, I. Time-to-Event Prediction with Neural Networks and Cox Regression. arXiv [stat.ML] (2019).

43. Kingma, D. P. & Ba, J. Adam: A Method for Stochastic Optimization. arXiv [cs.LG] (2014).

44. Buitinck, L. et al. API design for machine learning software: experiences from the scikit-learn project. arXiv [cs.LG] (2013).

45. Paszke, A. et al. PyTorch: An imperative style, high-performance deep learning library. Adv. Neural Inf. Process. Syst. 32, (2019).

46. Fey, M. & Lenssen, J. E. Fast Graph Representation Learning with PyTorch Geometric. arXiv [cs.LG] (2019).

47. McInnes, L., Healy, J. & Melville, J. UMAP: Uniform Manifold Approximation and Projection for Dimension Reduction. arXiv [stat.ML] (2018).

48. UMAP: Uniform Manifold Approximation and Projection for Dimension Reduction — umap 0.5 documentation. https://umap-learn.readthedocs.io/en/latest/.

